# The alternative initiation factor eIF2A regulates 40S subunit turnover in ribosome-associated quality control

**DOI:** 10.1101/2025.05.22.655611

**Authors:** Merve Yigit, Ramona Weber, Umesh Ghoshdastider, Anupam Das, Kivanc Nesanir, Fabiola Valdivia-Francia, Katie Hyams, David Taborsky, Peter F. Renz, Mark Ormiston, Clara Duré, Homare Yamahachi, Marko Jovanovic, Ataman Sendoel

## Abstract

The noncanonical translation initiation factor eIF2A plays critical roles in diverse cellular processes, including the integrated stress response, neurodegeneration and tumorigenesis. However, the precise molecular mechanism underlying eIF2A’s function remains poorly understood. Here, we exploit a TurboID-based proximity labeling combined with mass spectrometry to systematically map the interactome of eIF2A during homeostasis and stress. Combining polysome gradients with TurboID, we zoom into the interactions of eIF2A with the 40S small ribosomal subunit and map the eIF2A binding site close to the mRNA entry channel. We identify a network of interactors that link eIF2A to ribosome-associated quality control, including its strong interaction with G3BP1-USP10 complexes as well as RPS2 and RPS3. In the absence of eIF2A, RPS2 and RPS3 ubiquitination is diminished specifically upon ribosome stalling. 40S-specific footprinting in *eIF2A knockout* cells shows minimal changes in 5’UTR occupancy, consistent with a limited role for eIF2A in translation initiation. Using dynamic SILAC mass spectrometry, we characterize the novel function of eIF2A in ribosome-associated quality control and show that eIF2A antagonizes USP10-dependent rescue of 40S ribosomes, resulting in altered turnover of 40S subunits upon cellular stress. Collectively, our study identifies a previously unknown link between eIF2A and ribosome-associated quality control, implies that eIF2A promotes translation fidelity by tuning 40S ribosome rescue under stress and warrants further investigations into the role of ribosome-associated quality control in tumorigenesis.

## Introduction

Gene expression regulation is fundamental for maintaining cellular homeostasis and its deregulation is closely linked to the pathogenesis of various diseases including cancer (*1-3*). Within the gene expression cascade, mRNA translation enables rapid modulation of protein levels in response to a broad spectrum of intrinsic and extrinsic signals. This complex, multi-step process encompasses initiation, elongation, termination, and ribosome recycling, with initiation generally considered the rate-limiting step in protein production (*4-8*). The canonical translation initiation cascade involves the assembly of the ternary complex (TC) consisting of eukaryotic initiation factor 2 (eIF2), guanosine-5’-triphosphate (GTP) and the initiator methionyl-tRNA (Met-tRNA_?_), which delivers the initiator tRNA to the ribosomal P-site. Subsequently, the 43S preinitiation complex (PIC), comprising the 40S ribosomal subunit, the TC and additional initiation factors, is formed and recruited to the 5’end of the mRNA with the 7methylguanosine (m7G) cap binding complex eIF4F, forming the 48S initiation complex that scans the 5’untranslated region (UTR) to locate the start codon. Upon recognition of the start codon, GTP is hydrolyzed, followed by releasing initiation factors and joining of the 60S large ribosomal subunit to form the functional 80S ribosome (*9-12*).

A central mechanism in the control of translation initiation involves the inhibition of eIF2 by the integrated stress response (ISR). Cellular stressors such as amino acid starvation, endoplasmic reticulum (ER) stress or oxidative stress activate the ISR, leading to the phosphorylation of serine 51 on the alpha subunit of eIF2 (eIF2α) (*13-17*). Phosphorylation decreases the availability of active eIF2 and results in the repression of global translation. Nevertheless, certain stress response genes bypass global translational inhibition and remain translated, indicating the existence of alternative translation initiation mechanisms that are active under stress conditions (*18*).

Several alternative initiation factors, including eIF2D, MCTS-1/DENR, eIF5B and eIF2A, have been reported to interact with the 40S ribosome and facilitate eIF2-independent recruitment of Met-tRNA_?_ for transcript-specific translation initiation (*19-23*). eIF2D and MCTS-1/DENR have been shown to bind initiator tRNA and 40S ribosomes with high affinity (*24,25*). However, subsequent studies indicated that their primary role is in recycling empty tRNAs from 40S subunits stalled at termination codons, a mechanism likely central to their ability to modulate re-initiation downstream of upstream open reading frames (uORFs) (*20,21,23,26-29*). eIF5B promotes 60S ribosomal subunit joining and has similarly been reported to deliver initiator tRNA to the ribosome under stress conditions, particularly in the context of upstream open reading frames (uORFs) and internal ribosome entry sites (IRES) (*22,30-32*). Together, these observations point to broader functions for these noncanonical initiation factors beyond a direct role in translation initiation.

The alternative initiation factor eIF2A was initially identified in the 1970s and considered the functional analogue of prokaryotic IF2, but subsequent studies reassigned this role to the canonical initiation factor eIF2 (*4,33*). Unlike the heterotrimeric eIF2, eIF2A is a 65 kDa monomeric protein reported to act as a GTP-independent but codon-dependent Met-tRNA_i_ carrier for specific transcripts (*4,34*). While eIF2A is not essential for global translation initiation, it has been implicated in various aspects of alternative translation initiation, including IRES-dependent translation, uORF initiation, re-initiation following uORF translation, 40S sequestering, ribosomal frameshifting and initiation from near-cognate start sites, with sometimes contradictory findings (*13,33,35-40*).

Knockout models in *S. cerevisiae, C. elegans* and *M. musculus* are viable, indicating that eIF2A is not essential for cellular survival. However, its synthetic genetic interactions with eIF5B and eIF4E in yeast, with double deletions displaying severe slow growth phenotypes, suggest partial redundancy with these initiation factors (*34,41*). Furthermore, eIF2A has been implicated in the pathogenesis of various diseases. eIF2A knockout mice develop a metabolic syndrome characterized by fatty liver disease, obesity and insulin resistance, display reduced numbers of immune cells and have a shortened lifespan (*42,43*). Moreover, eIF2A has been associated with neurological diseases, such as myotonic dystrophy 2, familial amyotrophic lateral sclerosis and frontotemporal dementia (FTD), where eIF2A contributes to repeat-associated non-AUG (RAN) translation (*44,45*). Additionally, multiple lines of evidence implicate eIF2A in different aspects of tumorigenesis. In a squamous cell carcinoma allograft model, eIF2A was required for early stages of tumor formation, despite not affecting cellular proliferation or global translation *in vitro* (*46*), indicating a potential context-specific function in early oncogenic stress responses. Clinically, increased eIF2A levels correlate with shorter overall survival in head and neck squamous cell carcinoma patients (*46*). eIF2A has also been implicated in promoting cell survival upon chemotherapy (*47*), facilitating cellular adaptation through the integrated stress response (*13*) and supporting c-Src translation and cancer cell proliferation under stress (*48*).

Despite its involvement in the pathogenesis of various diseases, the precise molecular function of eIF2A remains unclear. Although eIF2A has been implicated in promoting non-AUG initiation under specific conditions, such as binding immunoglobulin protein (*BiP*) mRNA translation (*13*) in stress or the translation of N-terminally extended *PTEN* isoforms (*49*), recent ribosome profiling in eIF2A knockouts in yeast and human HeLa cells revealed minimal impact on global translation or uORF-mediated translational control (*50,51*). Altogether, these observations raise the possibility that eIF2A’s role may extend beyond a direct involvement in translation initiation, prompting us to systematically map its interactome and investigate the underlying molecular mechanisms.

Here, we employed TurboID-based proximity labeling coupled with mass spectrometry to define the interactome of eIF2A under homeostasis and stress. We identify a network associated with ribosome-associated quality control (RQC), including interaction with G3BP1-USP10 complexes and the 40S proteins RPS2 and RPS3. Loss of eIF2A diminishes RPS3 ubiquitination upon ribosome stalling and alters 40S turnover under stress, suggesting a surprising functional link between eIF2A and the rescue of 40S ribosomes during stress.

## Results

### TurboID-based proximity labeling to map the interactome of eIF2A

To identify eIF2A’s interaction partners and obtain insights into its role in translation, we employed a TurboID-based proximity labeling strategy combined with label-free mass spectrometry (*52*) (Fig. 1A). First, we generated N-terminal and C-terminal TurboID-fusion constructs for eIF2A (TurboID::eIF2A and eIF2A::TurboID), along with 2 control constructs (tGFP::TurboID and cytosolic TurboID) (Fig. 1B). Using low-titer lentiviral transduction, we introduced these TurboID constructs into *Hras*^*G12V*^; *Tgfbr2*^*-/-*^; *Eif2a*^*-/-*^ cells from a previously established allograft squamous cell carcinoma (SCC) model that depends on eIF2A for rapid tumor formation in *Nude* mice (*46,53*). We then selected stable lines expressing our TurboID constructs at levels comparable to endogenous eIF2A (Fig. S1A). The N-terminal TurboID::eIF2A construct partly rescued the subtle ribosome stoichiometry defects upon loss of eIF2A in polysome gradients, suggesting that the fusion constructs are functional (Fig. S1B). Next, we optimized biotinylation conditions in SCC cells expressing TurboID fusion constructs and determined that a minimum of 30 minutes of biotin incubation was necessary for robust proximity biotinylation, as confirmed by western blot analysis with streptavidin-HRP (Fig. S1C-E).

**Figure 1:**
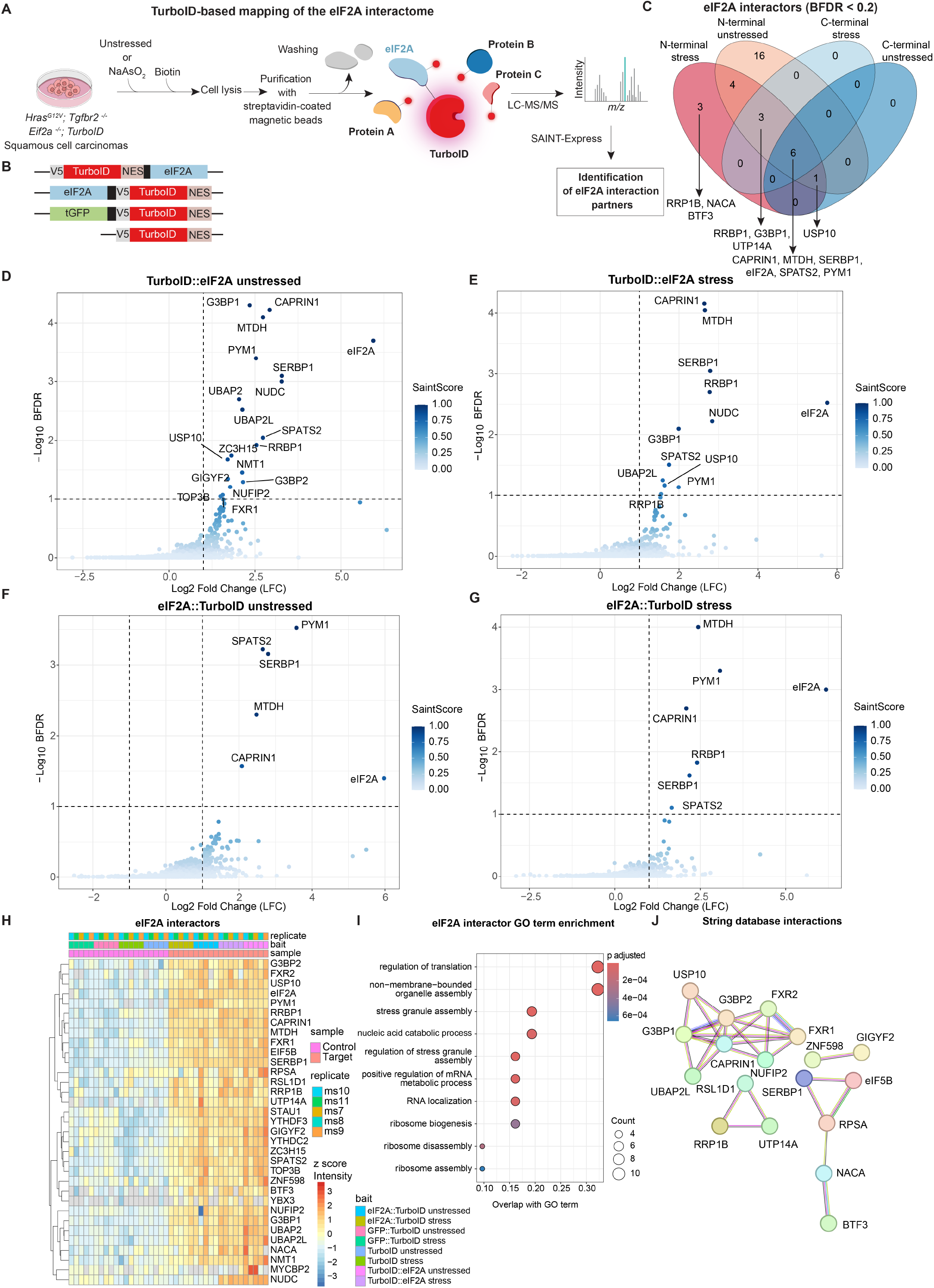
Mapping the interactome of eIF2A using TurboID-based proximity biotinylation. **(A)** Schematic outline of the TurboID-based proximity labelling of eIF2A interactors. *Hras*^*G12V*^; *Tgfbr2*^*-/-*^; *Eif2a*^*-/-*^ mouse squamous cell carcinoma cells expressing TurboID fusion constructs were subjected to sodium arsenite for 30 min or left untreated, followed by biotinylation (Methods). Cell lysates were purified with streptavidin-coated magnetic beads and were subjected to LC-MS/MS. eIF2A interactors were analyzed using MaxQuant and SAINTexpress. **(B)** Outline of the TurboID constructs that were used as baits for proximity labeling of eIF2A interactors. **(C)** Venn diagram showing the overlap of eIF2A interactors identified with N-terminal and C-terminal TurboID fusion constructs. Bayesian False Discovery Rate (BFDR) < 0.2. Candidate interactors were computed based on 5 independent experimental replicates. **(D-G)** Volcano plots displaying eIF2A interactors with Nterminal TurboID::eIF2A and C-terminal eIF2A::TurboID fusion constructs in unstressed and stress conditions. Interactors with BFDR < 0.1 are labeled. eIF2A is identified as the top enriched interactor (prey) due to autobiotinylation. Candidate interactors were computed based on 5 independent experimental replicates. **(H)** Heatmap displaying the intensity of top eIF2A interactors across the 5 replicates. **(I)** Gene Ontology (GO) enrichment of eIF2A interactors across all samples (BFDR < 0.2). **(J)** String database (STRING-db) highlights known interactions among the 33 eIF2A candidates (BDFR < 0.2). Disconnected nodes are not shown. Minimum required STRING-db interaction score: high confidence (0.7).

### eIF2A interacts with distinct groups of translational regulators

We carried out TurboID-based proximity labeling in SCC cells expressing either TurboID::eIF2A or eIF2A::TurboID fusion proteins, with tGFP::TurboID and cytosolic TurboIDexpressing cells as controls (Fig. 1A). In parallel, we also treated cells with sodium arsenite (NaAsO_2_) prior to biotin supplementation to induce the integrated stress response, resulting in phosphorylation of eIF2α and inhibition of canonical translation with reduced polysome fractions (Fig. S2A-B). We conducted 5 biological replicates of proximity labeling in unstressed and sodium arsenite-stress conditions for all 4 cell lines. After harvesting and purification of biotinylated proteins, samples were prepared for label-free mass spectrometry (LC-MS/MS) for protein identification. To identify eIF2A-specific interactors, we used MaxQuant and SAINTexpress, which calculates the probability of bait-prey interactions compared to controls (*54*). Principal component analysis (PCA) and heatmaps revealed that eIF2A-expressing cells were distinct from the TurboID controls in both unstressed and stress conditions (Fig. S3, S4A-B).

Using a Bayesian false discovery rate (BFDR) < 0.2, we identified 33 proteins as eIF2A interactors. 23 proteins specifically interacted with the N-terminal TurboID::eIF2A fusion protein, while 10 proteins were identified with both Nterminal and C-terminal constructs (Fig. 1C-G). Notably, 14 out of 33 proteins overlapped between stress and unstressed conditions, with only three interactors being stress-specific (Fig. 1C). Interestingly, we did not detect any C-terminus-specific interactors, suggesting that the N terminus of eIF2A is primarily responsible for protein-protein interactions (Fig. 1D-G, S5A). This notion is supported by structural insights as eIF2A’s N-terminal region contains an unconventional 9bladed β-propeller fold implicated in protein and RNA binding, while the C terminus is predicted to be intrinsically disordered (*55*).

As expected, Gene Ontology analysis revealed that eIF2A interactors are enriched in translational regulation, along with other biological processes such as stress granule assembly, RNA catabolic processes and ribosome biogenesis (Fig. 1I). Notably, STRING-db (*56*) analysis suggested that some of the eIF2A interactors form protein complexes linked to stress granule assembly and ribosome-associated quality control (RQC), such as the G3BP1, G3BP2, USP10 and CAPRIN1 (Fig. 1J, S5A-C), raising the possibility of a broader function of eIF2A within these complexes.

Using a stringent double cutoff of BFDR < 0.1 and SaintScore > 0.7, we identified a total of 17 high-confidence interactors, 10 of which were specific to the N-terminus. G3BP1, CAPRIN1 and USP10 were among the strongest interactors of eIF2A, irrespective of stress. G3BP1, CAPRIN1 and USP10 are key stress granule nucleating proteins responsible for the formation of these membrane-less organelles during stress (*57*). In addition, G3BP1 and USP10 are part of the deubiquitinase complex to rescue 40S ribosomes stalled in translation (*58*). Furthermore, eIF2A strongly interacted with SERBP1, a protein known to play a role in maintaining ribosome dormancy (*59,60*) and previously shown to interact with eIF2A (*61*).

Among the strong interactors, we also identified the two ubiquitin-associated proteins UBAP2 and UBAP2L (Fig. 1D-H). UBAP2 can trigger phase separation, as shown in the context of purinosome assembly (*62*). UBAP2 and UBAP2L may also regulate ubiquitination and degradation of cellular complexes such as RNA polymerase II following UVradiation (*63*). UBAP2L, on the other hand, binds to G3BP1 in stress granule assembly and fusion of UBAP2L to RNAtargeting CRISPR-Cas9 can strongly enhance global translation (*64-66*). Furthermore, another eIF2A interactor, PYM1, binds 40S ribosomes as well as exon-junction complexes (*67*). Finally, eIF2A interacted with several proteins associated with cancer-related processes such as MTDH, SPATS2 and RRBP1. MTDH (metadherin) has been widely implicated in cancer progression and acts as an RNA-binding protein to confer survival under stress (*68,69*). SPATS2 (spermatogenesis-associated serine rich 2), another RNA-binding protein, has been implicated in hepatocellular carcinoma and squamous cell carcinoma (*70,71*), whereas RRBP1, an ER-membrane protein that mediates mRNA localization and modulates ER stress, is associated with increased aggressiveness across several tumor types (*72,73*).

To confirm the TurboID-based proximity labeling interactions, we used orthogonal methods and performed co-immunopurification of endogenous eIF2A with G3BP1 or SERBP1 in SCC cells. To rule out cell-type-specific effects, we additionally validated our findings in HEK 293T cells throughout this study. eIF2A interacted with G3BP1 and SERBP1 in wild-type cells under both basal and stress conditions (Fig. 2A). Given that G3BP1 is a key protein involved in stress granule formation, we further investigated the cellular localization of eIF2A under stress. eIF2A did not affect stress granules formation, as *eIF2A knockout* cells retained the ability to form G3BP1 foci under stress (Fig. 2B-C). We observed partial co-localization of endogenous eIF2A with G3BP1 in unstressed cells; however, although eIF2A was present in stress granules upon stress, its signal did not specifically enrich in stress granules but also remained cytoplasmic (Fig. 2B-C, S6A-B). Furthermore, eIF2A did not exclusively localize to stress granules even after prolonged stress exposure (Fig. 2C). Together, these findings validate the eIF2A interactions identified in our TurboID experiments and suggest that eIF2A may contribute to translational regulation beyond its assumed role in translation initiation.

**Figure 2:**
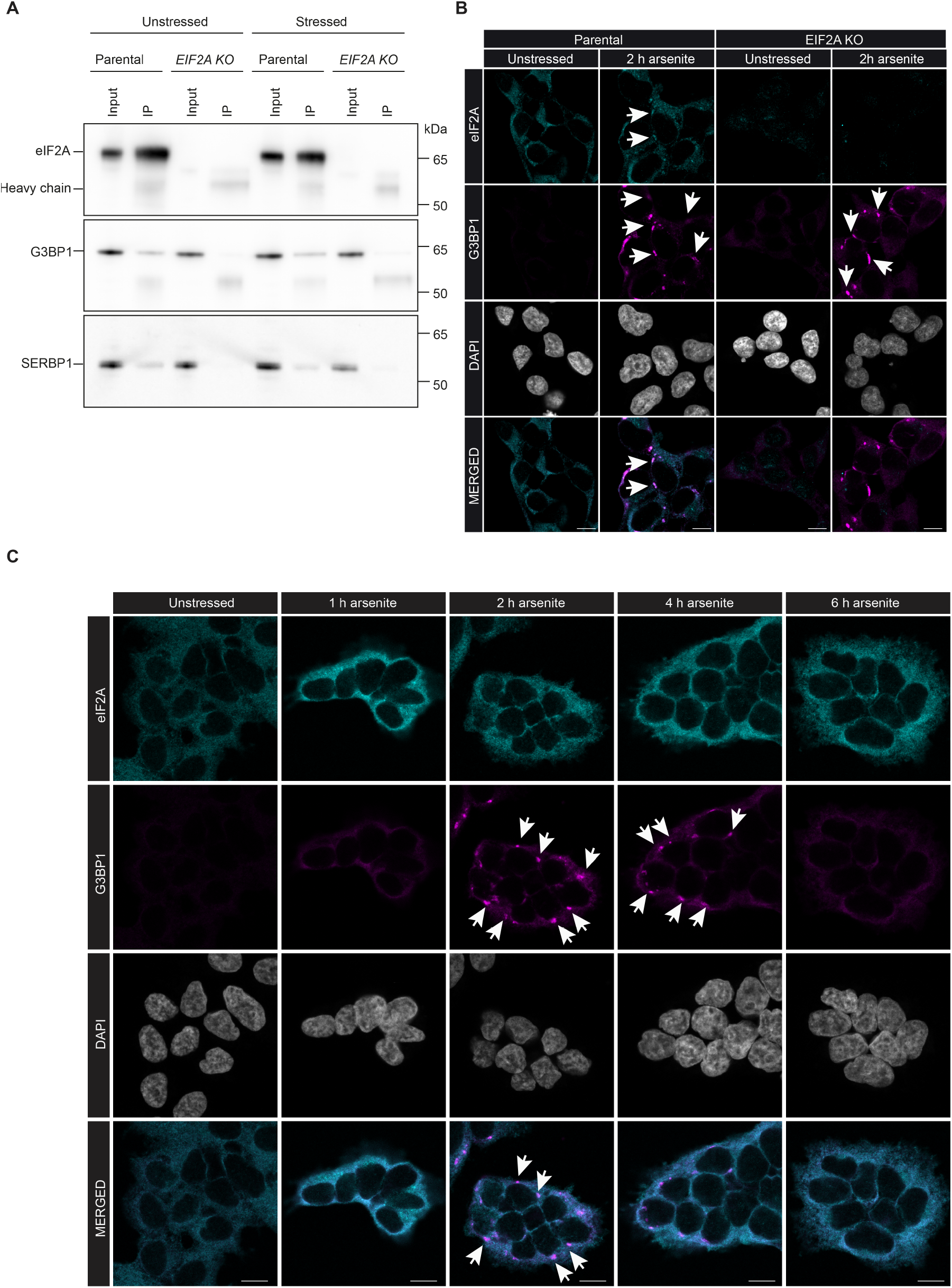
Validation of eIF2A interactors. **(A)** Co-immunopurification of eIF2A interactors G3BP1 and SERBP1 in control and sodium arsenite-treated HEK 293T control and *eIF2A knockout* cells. Parental and *eIF2A knockout* cells were stressed with 50 µM sodium arsenite for 2 hours. **(B)** Confocal microscopy images displaying eIF2A localization in HEK 293T cells. Parental and *eIF2A knockout* cells were stressed with 500 µM sodium arsenite for 2 hours and co-stained with the stress granule marker, and eIF2A interactor, G3BP1. Scale bars, 10 µm. **(C)** Confocal microscopy of eIF2A co-localization with G3BP1 in a time course upon sodium arsenite stress. Despite co-localization with cytosolic G3BP1, eIF2A did not exclusively localize to stress granules upon arsenite treatment. Scale bars, 10 µm.

### Selective 40S ribosomal proximity labeling reveals interaction with RQC factors

Several studies have reported interactions between eIF2A and the 40S ribosomal subunit, including evidence from *in vitro* translation assays (*40*). Thus, we were surprised that we did not detect any ribosomal proteins or initiation factors as interactors of eIF2A in our proximity biotinylation experiments (Fig. 1D-H). We surmised that the discrepancy might reflect the TurboID approach’s reduced sensitivity to capture transient interactions. Additionally, because TurboID preferentially labels proteins in close proximity, eIF2A might associate indirectly with the 40S ribosome within larger complexes, potentially reducing labeling efficiency due to longer spatial distances. Consistent with the possibility of transient 40S interactions, eIF2A predominantly localized to the free ribonucleoprotein (RNP) fraction in polysome gradients in SCC cells, with lower association to the 40S fraction (Fig. 3B). This implies that interactions identified by our TurboID approach may primarily originate from the free RNP complexes, potentially missing more transient associations with the 40S ribosome. To investigate potential eIF2A interactions with the 40S ribosome, we performed selective 40S-TurboID proximity labeling combined with label-free mass spectrometry in our TurboID::eIF2A and tGFP::TurboID-expressing cells (Fig. 3A). Following biotinylation, we isolated 40S and 80S (both monosomes and polysomes) ribosomal fractions using sucrose density gradient separation, enabling the identification of eIF2A-associated interactors with the ribosome (Fig. 3C-H, S4C-E).

**Figure 3:**
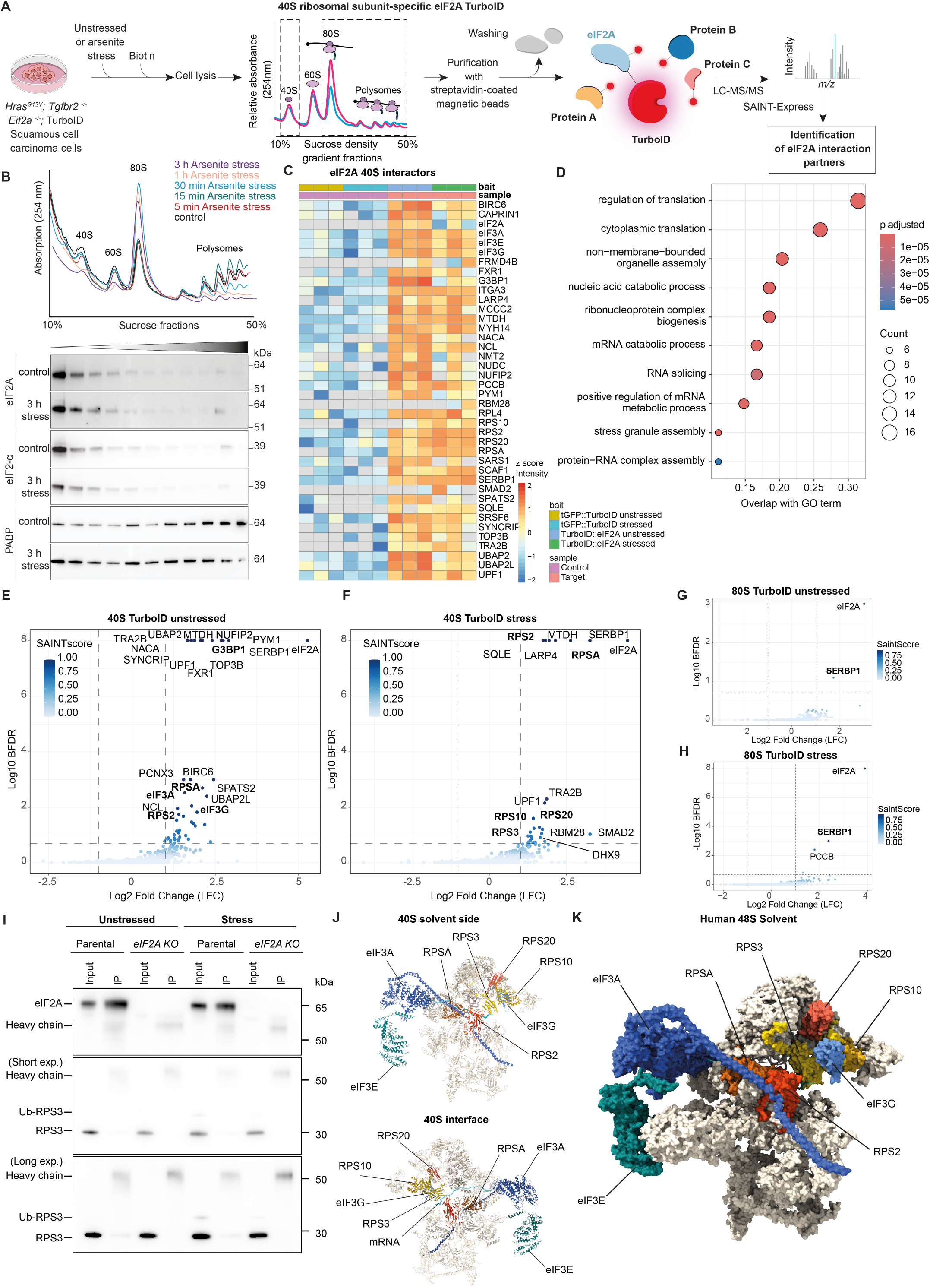
40S-specific proximity biotinylation uncovers interactors associated with RQC. **(A)** Schematic outline of 40S TurboID proximity labeling approach. Cells were stressed with sodium arsenite or left untreated, followed by biotinylation. The lysates were subjected to sucrose density gradient fractionation followed by streptavidin-coated magnetic beads purification and LC-MS/MS. tGFP::TurboID was used as control. Candidates were analyzed using FragPipe and SAINTexpress software. **(B)** Sucrose density gradients of HEK 293T cells were collected over a time course following sodium arsenite-induced stress. Western blot images of polysome fractions show eIF2A and the canonical initiation factor eIF2α associated with free RNPs and 40S subunits. **(C)** Heatmap displaying the intensity of top eIF2A interactors across the 3 independent replicates (BFDR < 0.1). **(D)** Gene Ontology enrichment of top 40S-specific eIF2A interactors across all conditions (BFDR < 0.2). **(E-H)** Volcano plots displaying eIF2A interactors on the 40S and 80S ribosomes. Candidate interactors were computed based on 3 independent experimental replicates using FragPipe and SAINTexpress software. **(I)** Co-immunopurification of the eIF2A interactor RPS3 in control and sodium arsenite-stressed HEK 293T cells. In addition, stress resulted in ubiquitination of RPS3 in input, which was reduced in *eIF2A knockout* cells. Of note, the co-immunopurifications presented in Fig. 2A and 3I were probed with different antibodies in the same experiment and therefore the same eIF2A blot is shown in both panels. **(J)** Cryo-EM structure of the mammalian cytosolic 40S ribosome (*76*) (Homo sapiens, 3.70 Å) on the solvent side (upper panel) and interface side (lower panel). eIF2A interactors are labeled and located close to the mRNA entry channel. **(K)** Structure of the human 48S initiation complex (*76*) highlighting the eIF2A interactors RPS2, RPS3, RPS10, RPS20, RPSA as well as eIF3A, eIF3E and eIF3G.

In the 80S-TurboID proximity labeling, SERBP1 emerged as the single significant interactor in both unstressed and stress conditions, underscoring a robust association with this dormant ribosome factor (Fig. 3G-H). In contrast, assessing the interactions in the 40S fraction, we identified a total of 40 eIF2A interactors (BFDR < 0.1) in unstressed and stress conditions (Fig. 3C, 3E-F). Of those, 33 were among the strong interactors (SAINTScore > 0.7). First, as in the classic TurboID approach, we observed a strong interaction with G3BP1, CAPRIN1 and SERBP1. Notably, with the 40STurboID approach, we also identified 48S ribosome-specific eIF2A interactions. These included RPS2, RPS3, RPS10, RPS20 and RPSA, as well as the eukaryotic initiation factor 3 subunits eIF3A, eIF3E, and eIF3G (Fig. 3C, 3E-F, S4D). We selected RPS3 for further validation using orthogonal assays and confirmed its robust association with eIF2A by coimmunopurification (Fig. 3I).

These results were interesting for several reasons. First, RPS2, RPS3, RPS10, RPS20, as well as initiation factors eIF3A, eIF3E, and eIF3G, are all located close to the mRNA entry channel of the 40S ribosomal subunit (Fig. 3J-K) (*74-76*). Thus, our data imply that the N terminus of eIF2A may interact with the 40S ribosome close to the mRNA entry channel. This notion is consistent with recent enhanced crosslinking and immunoprecipitation (eCLIP) data showing that eIF2A robustly interacts with regions of the 18S rRNA between helices 21es6B and 21es6C, situated near the mRNA entry channel (*38*). Second, RPS2, RPS3, RPS10 and RPS20 are well-established targets of regulatory ubiquitination during the ribosome-associated quality control (RQC) pathway following ribosome stalling and collision (*77*), prompting us to consider whether eIF2A participates in RQC. Of note, in co-immunopurification experiments, we identified a ubiquitinated form of RPS3 under stress, indicating activation of the RQC pathway (Fig. 3I). Interestingly, stress-induced RPS3 ubiquitination was markedly reduced in *eIF2A knockout* cells compared to parental controls (∼85% reduction in Ub-RPS3:RPS3 ratio), suggesting a possible defect in the RQC pathway upon stress (Fig. 3I, input lane). Third, two of the key eIF2A interactors, G3BP1 and USP10, are required for the deubiquitination of RPS2, RPS3 and RPS10 to rescue ubiquitinated 40S subunits stalled in translation (*58*).

Given that these observations hinted at a role beyond translation initiation, we first set out to formally exclude a contribution of eIF2A to initiation in our system. Recent ribosome profiling experiments in yeast and human cells supported the notion that eIF2A plays only a minor role in translation initiation, with relatively few transcripts showing altered translation profiles (*50,51*). To further assess whether eIF2A contributes to scanning and translation initiation in our system, we performed 40Sspecific ribosome profiling using translation complex profiling (TCP-seq) in parental and *eIF2A knockout* SCC cells (*78,79*). The 40S metagene profiles of parental and *eIF2A knockout* SCC cells were largely indistinguishable in unstressed and stress conditions (Fig. 4A-B). Furthermore, differential 40S footprint analysis in the 5’UTR revealed only a small number of transcripts with differential reads in the 5’UTR, excluding the arguably best-studied uORF-regulated gene *Atf4*, which showed no significant changes (Fig. 4C-D). Moreover, only 2 transcripts showed differential footprints in the 5’UTR when comparing stress versus unstressed conditions (Fig. 4C). Of note, proteomics analysis in *eIF2A knockout* SCC cells revealed also only a modest number of differentially expressed proteins in homeostasis (Table S1).

**Figure 4:**
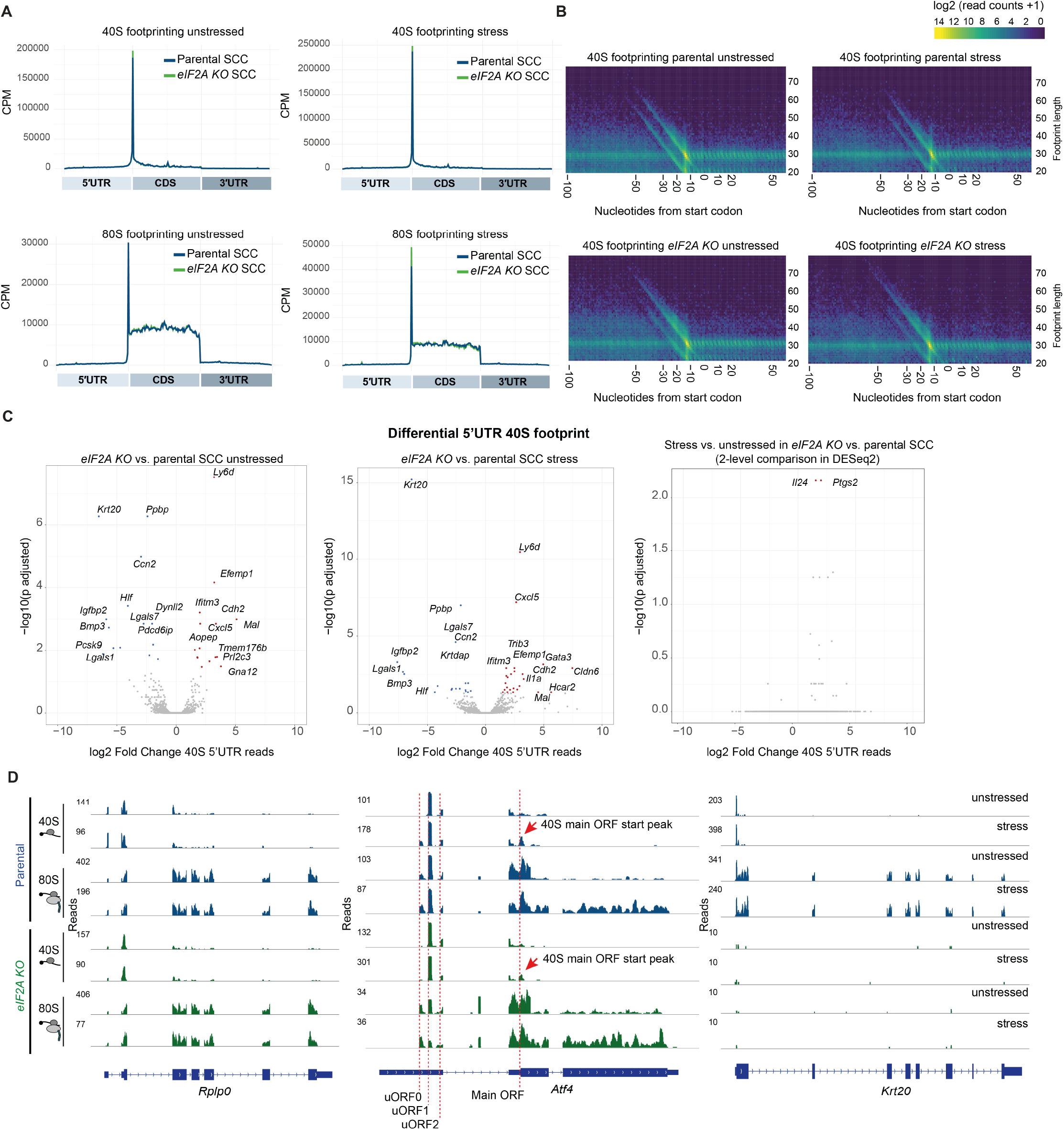
40S-specific ribosome profiling using TCP-seq. **(A**) Metagene profiles comparing 40S-specific translation complex profile sequencing (TCP-seq) and conventional (80S) ribosome profiling in parental and *eIF2A knockout* SCC cells. CPM, counts per million. **(B)** 40S footprint distribution in parental and *eIF2A knockout* SCC cells. **(C)** Differential 5’UTR footprint distribution analysis in parental and *eIF2A knockout* SCC cells under unstressed and sodium arsenite (50 µM, 2 hours) stress conditions. Differential footprint analysis was conducted using DESeq2 with two independent replicates per condition. No transcript-level normalization was applied. **(D)** Representative footprint tracks for the housekeeping gene *Rplp0*, the stress-responsive gene *Atf4* and *Krt20*, one of the few transcripts showing altered footprint occupancy in the 5’UTR. Blue tracks, control; green tracks, *eIF2A knockout* SCC cells.

In summary, these data support the view that eIF2A plays a minor role in translation initiation but may instead be involved in the RQC pathway, potentially regulating the deubiquitination of stalled 40S ribosomes through its association with G3BP1-USP10 complexes.

### eIF2A impacts ribosome-associated quality control during stress

To assess whether eIF2A is involved in the RQC pathway, we performed collision and stalling assays by treating cells with either sodium arsenite or anisomycin. Anisomycin induces ribosome collisions at intermediate doses and ribosome stalling at higher doses (*80*). Since the RQC pathway has been extensively studied in HEK 293T cells, we included both SCCs and HEK 293T cells in our experiments to validate these observations across cell types (Fig. S7F). Following stress, we monitored RPS3 ubiquitination by western blotting as a marker of RQC activation. As expected, RPS3 was ubiquitinated in parental cells in response to both intermediate and high doses of anisomycin, as well as upon sodium arsenite treatment (Fig. 5A). Notably, *eIF2A knockout* cells showed a pronounced reduction in RPS3 ubiquitination across all stress conditions compared to parental cells, as quantified from three independent replicates in both SCC and HEK 293T cells (Fig. 5A-B, S7A-E). This result aligned with the reduced RPS3 ubiquitination observed in our co-immunopurification blots (Fig. 3I). The reduction was most prominent following highdose anisomycin treatment, with HEK 293T *eIF2A knockout* cells showing virtually no increase in RPS3 ubiquitination upon stalling (Fig. 5A-B), indicating that eIF2A may specifically impact RPS3 ubiquitination upon ribosome stalling rather than following ribosome collision. Furthermore, employing sucrose density gradient fractionation in high-dose anisomycin and arsenite-treated cells showed that the ubiquitinated RPS3 is mainly found in the 40S fraction, with little RPS3 ubiquitination in the polysome fractions. As expected, *eIF2A knockout* cells showed a marked reduction of ubiquitinated RPS3 specifically in the 40S fraction upon ribosome stalling (Fig. 5C). Moreover, we observed a similar reduction in RPS2 ubiquitination in *eIF2A knockout* cells, which was also most pronounced upon high-dose anisomycin-dependent ribosome stalling (Fig. S8A-B). Our findings therefore pointed to a model in which eIF2A interacts with G3BP1– USP10 complexes to either promote the ubiquitination of RPS2 and RPS3 or antagonize their deubiquitination by USP10.

**Figure 5:**
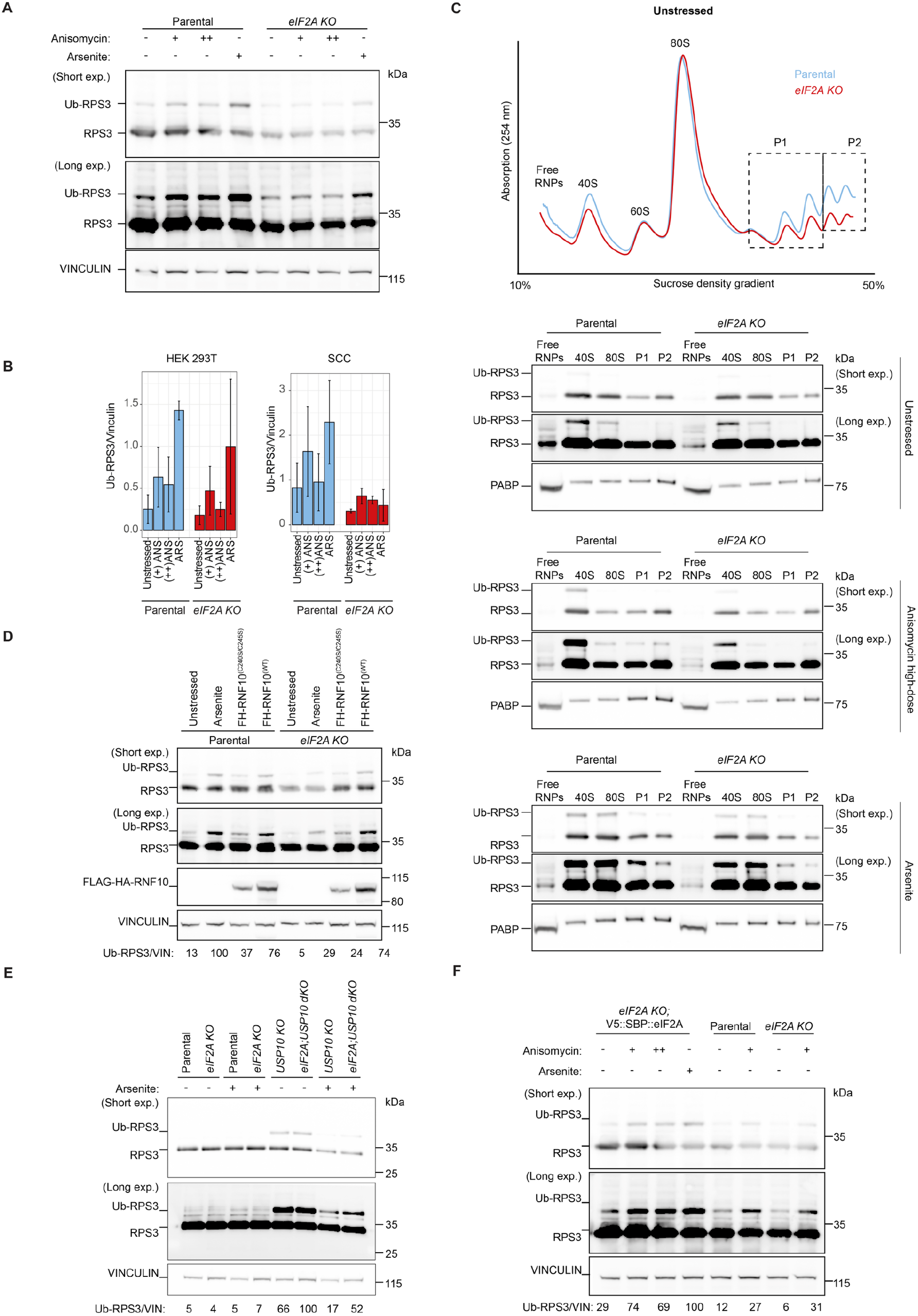
Loss of eIF2A attenuates RPS3 ubiquitination upon stress. **(A)** Western blot analysis displaying RPS3 ubiquitination upon anisomycin-induced ribosome collision (+) or stalling (++). HEK 293T cells were incubated with intermediate dose (+, 1 µM) or high dose (++, 100 µM) anisomycin for 30 minutes prior to harvesting. In parallel, cells were stressed with sodium arsenite for 2 hours. Short exp.: shorter exposure. Long exp.: longer exposure. **(B)** Quantification of RPS3 ubiquitination bands in HEK 293T cells and SCCs shown in Figure 5A and Figure S7 (n=3 each). **(C)** Anisomycin assay sucrose density gradients displaying polysome distribution in parental and *eIF2A knockout* HEK 293T cells upon anisomycin-induced ribosome stalling. Western blot images of the different fractions show RPS3 ubiquitination mainly in the 40S fraction. RNP, ribonucleoproteins. **(D)** Western blot images displaying overexpression of human RNF10 constructs (FLAG::HA::RNF10^Wildtype^ and FLAG::HA::RNF10^(C240S/C245S)^ catalytically inactive mutant) in parental and *eIF2A knockout* HEK 293T cells. **(E)** Western blot images displaying RPS3 ubiquitination upon loss of USP10 in SCCs. *Usp10 knockout* cells show increased RPS3 ubiquitination in the absence of stress. *eIF2A; Usp10* double knockout cells showed no significant change in RPS3 ubiquitination levels relative to *Usp10* single knockout cells. As previously reported (*83*), arsenite treatment results in reduced RPS3 ubiquitination. **(F)** eIF2A expression in *eIF2A knockout* cells markedly increases RPS3 ubiquitination. eIF2A expression was restored in *eIF2A knockout* cells by lentiviral infection with a V5::SBP::eIF2A construct. Cells were treated with intermediate-dose (+) or high-dose (++) anisomycin for 30 minutes as well as sodium arsenite for 2 hours.

We therefore sought to systematically investigate the potential involvement of eIF2A in the different arms of the RQC pathway. First, we assessed ZNF598-dependent ribosome collision sensing, one of eIF2A’s low-confidence interactors (BFDR < 0.2 and SAINTScore < 0.7), which detects ribosome collisions and triggers the downstream RQC response (*81*). We employed collision-inducing luciferase reporters, which previously led to rapid transcript degradation in wild-type but not in *ZNF598 knockout* cells (*58*). However, in contrast to the loss of ZNF598, *eIF2A knockout* HEK 293T cells failed to increase the expression of the collision-inducing luciferase reporter, suggesting that eIF2A functions in the RQC pathway through a mechanism distinct from ZNF598-mediated collision sensing (Fig. S8C). This observation was consistent with our finding that eIF2A may primarily influence RQC in response to ribosome stalling rather than collision. Second, we focused on RNF10, which ubiquitinates RPS2 and RPS3 upon stalling of elongating ribosomes or scanning 40S ribosomes in the context of initiation RQC (iRQC) (*82-84*). Since the mechanism of RNF10 recruitment to the 40S ribosome remains unknown, we asked whether eIF2A could mediate its recruitment to stalled ribosomes. To test this hypothesis, we used a doxycycline-inducible FLAG-HA tagged RNF10 overexpression construct (*82*). As expected, overexpression of RNF10 in HEK 293T cells increased ubiquitinated RPS3 levels (*82*). However, we observed no apparent difference between parental and *eIF2A knockout* cells, suggesting that eIF2A acts either downstream or in parallel of RNF10mediated RPS3 ubiquitination (Fig. 5D, S8D).

Third, we focused on the two key eIF2A interactors, G3BP1 and USP10, which act downstream of RNF10dependent 40S subunit ubiquitination. While G3BP1 serves as a scaffold, USP10 acts as the deubiquitinase for RPS2 and RPS3 to rescue stalled 40S subunits from degradation (*58,82-84*). Given the close interaction of eIF2A with USP10, we considered two possible scenarios in which eIF2A either directly or indirectly antagonizes USP10 function. In the first scenario, eIF2A acts directly on USP10 to antagonize RPS3 deubiquitination upon stalling, in which case double *USP10; eIF2A knockout* cells would not reduce the *USP10 knockout* phenotype (as its function would depend on USP10). Or eIF2A indirectly antagonizes USP10-dependent RPS3 deubiquitination, in which case double *USP10; eIF2A knockout* cells should show reduced RPS3 ubiquitination compared to USP10 mutants (similar to RNF10; USP10 double mutants showing strongly reduced RPS3 ubiquitination (*82*)). We therefore generated CRISPR-mediated *Usp10; eIF2A double knockout* SCC cells (Fig. S8E-F). While the loss of USP10 resulted – as expected – in increased RPS3 ubiquitination, we observed no obvious reduction between parental and *eIF2A knockout* cells (Fig. 5E, S8E). Consistent with previous findings (*83*), sodium arsenite reduced RPS3 ubiquitination in *Usp10 knockout* cells as well as in *Usp10; eIF2A double knockout* SCCs (Fig. 5E, S8E). These findings support a working model wherein eIF2A may directly oppose USP10-mediated deubiquitination of RPS3.

Fourth, we tested a potential preferential involvement of eIF2A in iRQC. *eIF2A knockout* cells displayed, comparable to anisomycin treatment, a 50-70% reduction in ubiquitinated RPS3 following the translation initiation inhibitor harringtonine (Fig. S9A-B), suggesting that eIF2A also impacts stalled ribosomes during iRQC. Finally, to confirm eIF2A-dependent changes in RPS3 ubiquitination upon ribosome stalling, we stably expressed eIF2A by lentiviral transduction of a V5-tagged eIF2A construct and monitored RPS3 ubiquitination in unstressed and anisomycin-treated conditions. As expected, RPS3 ubiquitination upon anisomycin was reduced in *eIF2A knockout* HEK 293T cells (Fig. 5F). Interestingly, rescuing eIF2A expression in unstressed *eIF2A knockout* cells was already sufficient to increase RPS3 ubiquitination, suggesting that eIF2A re-expression may be sufficient to block USP10-mediated deubiquitination of low levels of physiological ribosome stalling events (Fig. 5F). Furthermore, upon anisomycin treatment, eIF2A rescue showed clearly increased RPS3 ubiquitination compared to parental or *eIF2A knockout* cells (Fig. 5F). Collectively, this systematic investigation, together with our TurboID data, suggest a working model wherein eIF2A may closely interact with G3BP1-USP10 complexes to antagonize USP10-dependent RPS3 deubiquitination upon ribosome stalling.

### eIF2A facilitates faster ribosomal protein turnover

The balance between RNF10 and USP10 during ribosome stalling plays a critical role in maintaining translation dynamics. RNF10 flags ribosomes stalled during initiation or elongation, which – if unresolved – triggers 40S subunit degradation (*58,82*). However, in homeostasis, the excess of USP10 over RNF10 favors the deubiquitination and rescue of 40S subunits (*83*), preventing energetically costly degradation of ribosomal subunits. *USP10 knockout* or RNF10 overexpressing cells enhance 40S ribosomal degradation and turnover rates (*58,83*). Thus, our working model would predict that *eIF2A knockout* cells shift the balance toward slower turnover of 40S subunits upon stalling.

To systematically examine whether eIF2A impacts ribosomal protein turnover rates, we performed dynamic SILAC (stable isotope labeling by amino acids in cell culture) mass spectrometry in SCC and HEK 293T cells (Fig. 6A) (*85*). Parental and *eIF2A knockout* cells were grown in light isotope-labeled media. We then switched to heavy isotopelabeled media along with low-dose sodium arsenite stress in a time course over 48 hours, allowing us to track protein turnover of newly synthesized (heavy isotope labeled) compared to degraded (light isotope labeled) proteins by LC-MS/MS (*85*). We found that *eIF2A knockout* cells did not majorly alter global protein turnover, as highlighted by comparable heavy:light ratios at the 24-hour time point or even slightly increased turnover at earlier time intervals (Fig. 6B, 6E). In contrast, within the cohort of 40S ribosomal proteins, *eIF2A knockout* SCCs showed strongly reduced heavy:light ratios compared to parental cells, suggesting a reduced protein turnover (Fig. 6C, 6F). This effect was specific to 40S ribosomal proteins, with no significant changes observed in other related protein groups, such as eukaryotic initiation factors (Fig. 6D, 6G, S10A-C). We confirmed these findings using median normalization across both dynamic SILAC replicates and similarly observed significantly reduced 40S turnover rates when combining the two dynamic SILAC datasets (Fig. 6H). In agreement with these observations, the 40S ribosomal protein half-lives were significantly increased in *eIF2A knockout* SCCs, further supporting reduced protein turnover (Fig. 6I).

**Figure 6:**
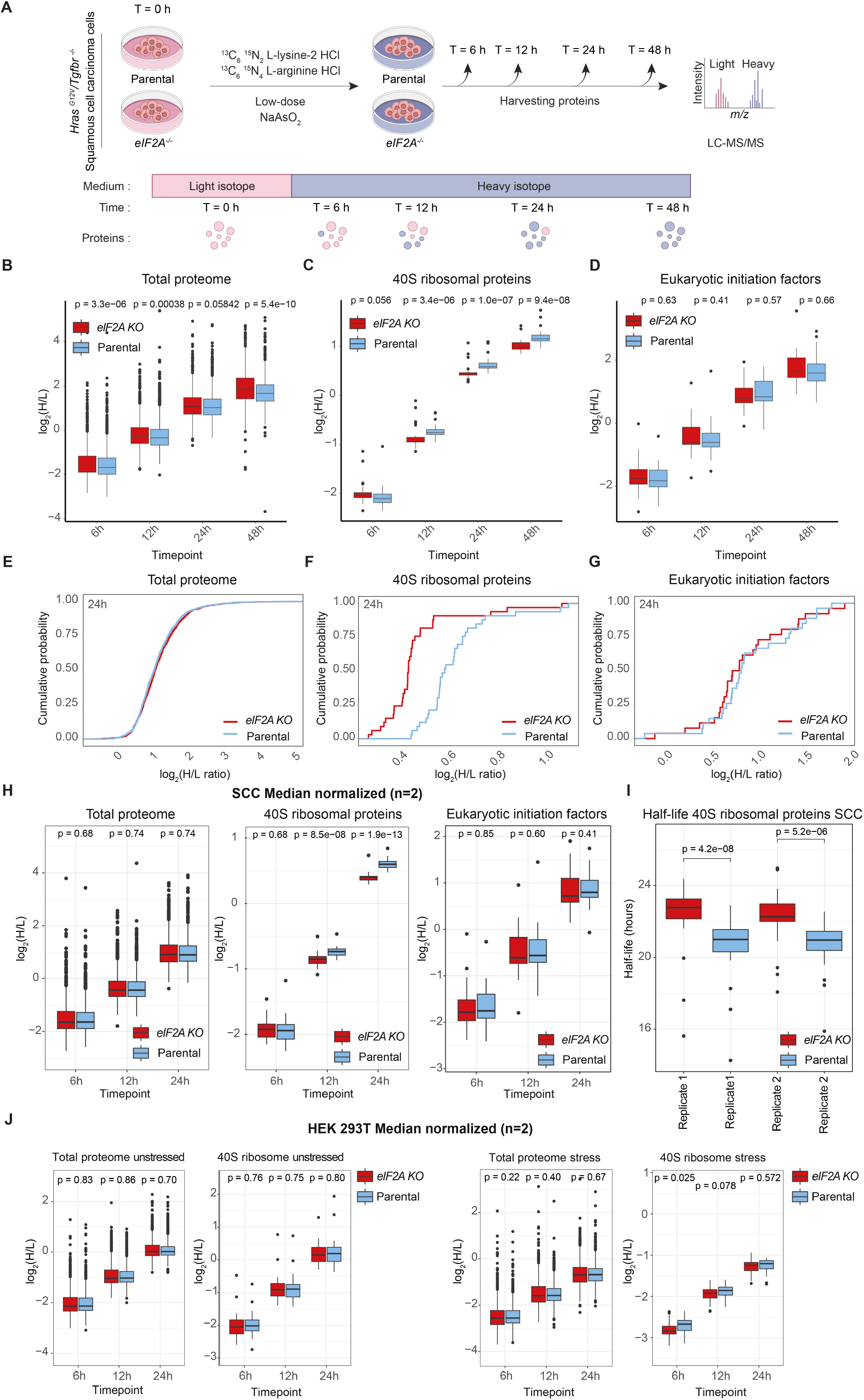
Loss of eIF2A reduces 40S subunit turnover upon stress. **(A)** Schematic outline of stable isotope labeling of amino acids in cell culture (SILAC) LC-MS/MS experiment. SCCs were kept in the light isotopecontaining medium. At t = 0 h, cells were switched to a media containing low-dose sodium arsenite alongside a heavy isotope-containing medium. Proteins were collected at 6 h, 12 h, 24 h and 48 h time points and subjected to LC-MS/MS in 2 independent replicate runs. Proteomics analysis was done using MaxQuant. **(B-D)** *eIF2A knockout* cells have reduced heavy:light (H/L) ratio in the 40S ribosomal proteins. Boxplots displaying H/L ratio of SCC cells over several time points in the total proteome **(B)**, 40S small ribosomal proteins **(C)** and eukaryotic initiation factors **(D). (E-G)** Cumulative distribution of H/L ratio of proteins in parental and *eIF2A knockout* SCCs at the 24-hour time point in the total proteome **(E)**, 40S small ribosomal proteins **(F)** and eukaryotic initiation factors **(G). (H)** Boxplots showing the median-normalized and combined H/L ratio of SCCs over the different time points in SCC cells. For each time point, each sample was median-normalized and averaged across two replicates. **(I)** 40S ribosomal half-life is significantly longer in *eIF2A knockout* SCCs. L fraction for each protein was calculated from H/L ratios using the formula L/(H+L) = 1/(1+H/L). The resulting L fraction values were then fitted to an exponential decay function L/(H+L) = exp(-kt), where k is the rate constant and t is time. Curve fitting was performed using the curve_fit function from the scipy Python library, requiring a minimum of three time points with complete data for each protein. Finally, protein half-lives were determined as ln(2)/k. **(J)** Boxplots showing the median-normalized and combined H/L ratio of unstressed and lowdose arsenite (5 µM) stressed HEK 293T over a time course of 24 hours. Each sample and time point was median-normalized and averaged across two replicates.

To extend our findings, we performed similar dynamic SILAC experiments in HEK 293T cells. An advantage of using HEK 293T was that it enables a steady-state condition upon medium switch from light to heavy, which was not achievable in SCC cells due to differing medium requirements (hence, no unstressed condition could be achieved in SCC). We therefore applied unstressed and lowdose sodium arsenite conditions over a 24-hour time course. In agreement with the SCC data, although HEK 293T protein turnover showed higher variance, 40S ribosomal turnover was also reduced in *eIF2A knockout* cells in stress but not in homeostasis, indicating that eIF2A specifically impacts 40S turnover in stress (Fig. 6J, S10D-E). Collectively, these findings in SCC and HEK 293T cells suggest that eIF2A specifically facilitates 40S subunit turnover under stress.

Finally, given that higher eIF2A transcript levels were associated with shorter overall survival in head and neck squamous cell carcinomas (*46*), we investigated whether ubiquitination of small ribosomal proteins in the context of RQC was generally altered in cancer patients. Among the available proteogenomic cancer datasets, only the one in lung squamous cell carcinoma included data on the ubiquitylproteome (*86*). Comparing specifically the ubiquitylproteome of tumor and healthy tissues, we found that RPS2, RPS3 and RPS10 but not RPS20 showed significantly higher overall ubiquitination levels (Fig. 7A). Given that lysine 214 (K214) on RPS3 is the crucial residue of monoubiquitination during RQC (*58*), we next examined site-specific changes. Interestingly, while the majority of residues showed no difference, ubiquitination at RPS3 K214 was markedly increased in cancer samples (Fig. 7B). Furthermore, increased RNF10 levels, which may enhance the capacity to flag stalled ribosomes similar to eIF2A, correlated with shorter overall survival in head and neck squamous cell carcinoma (Fig. 7C). These findings hint at a potentially broader involvement of RQC-dependent 40S turnover in the context of cancer progression and warrant further exploration of the role of ribosome-associated quality control in tumorigenesis.

**Figure 7:**
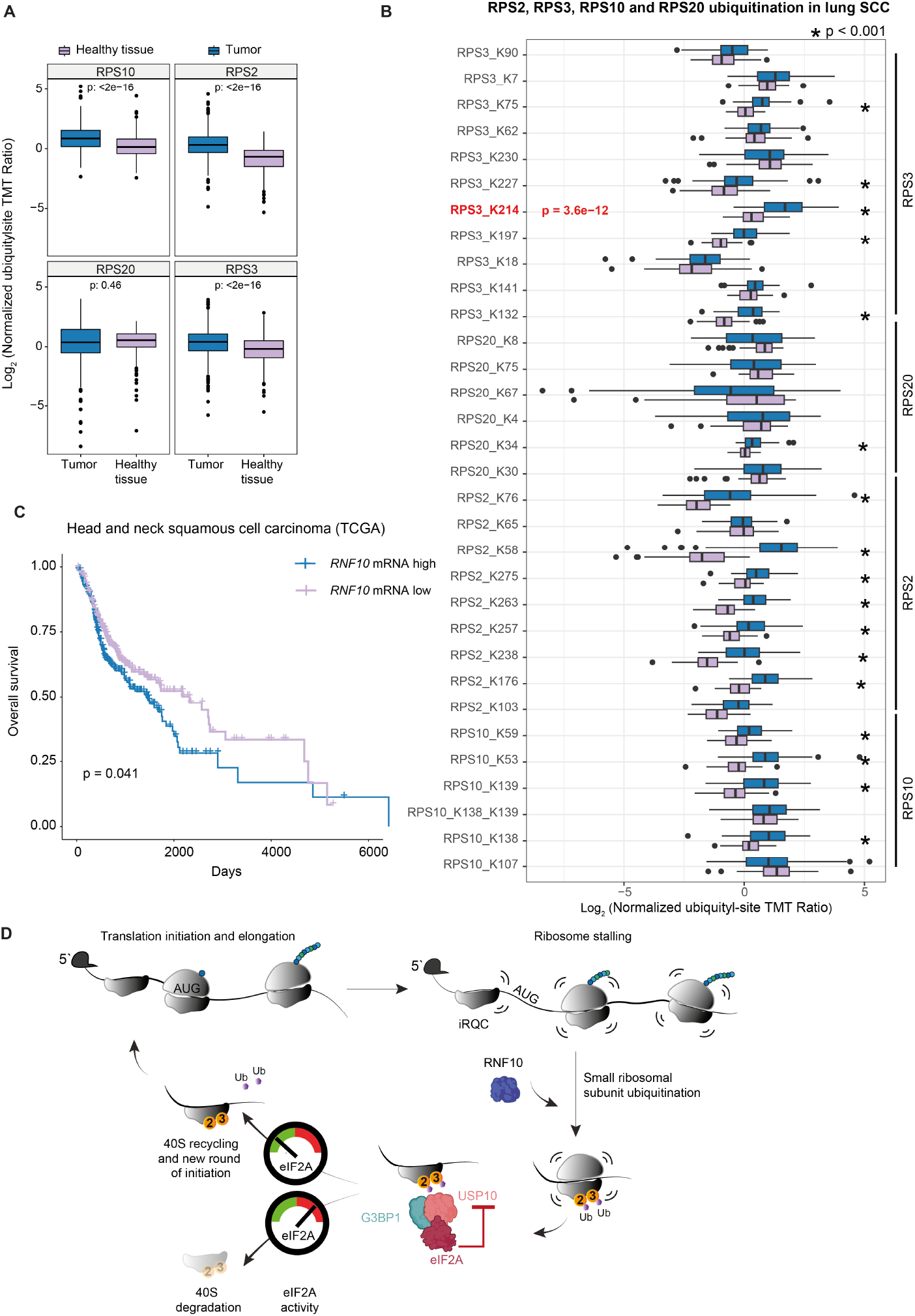
RPS3 is strongly ubiquitinated at the K214 residue in lung SCC patients. **(A)** Boxplots display the ubiquitination of small ribosomal proteins RPS2, RPS3, RPS10 and RPS20 in tumor and adjacent healthy tissue.**(B)** Boxplots showing site-specific ubiquitination in 40S ribosomal proteins RPS2, RPS3, RPS10 and RPS20. **(C)** Higher RNF10 mRNA levels correlate with shorter overall survival in head and neck squamous cell carcinoma patients (TCGA). **(D)** The proposed working model in which eIF2A binds to G3BP1-USP10 complexes to antagonize USP10-mediated deubiquitination and facilitate 40S subunit degradation during ribosome-associated quality control.

## Discussion

Although discovered over five decades ago and implicated in various alternative initiation mechanisms and diseases, the molecular function of eIF2A has remained unclear. Here, we employed an unbiased TurboID-based proximity biotinylation strategy to map the interactome of eIF2A in homeostasis and stress. We identify a surprising functional association between the alternative initiation factor eIF2A and 40S subunit turnover in the context of ribosome-associated quality control (Fig. 7D).

Protein production is one of the most energetically expensive processes in the cell. Eukaryotic cells have developed surveillance mechanisms that recognize and resolve ribosomes stalled on defective mRNAs, thereby maintaining translation fidelity. While collided ribosomes are marked by ZNF598, the E3 ubiquitin ligase RNF10 initiates site-specific monoubiquitination of RPS2 and RPS3upon stalling to flag 40S subunits for programmed degradation (*82,84*). G3BP1-USP10 complexes can rescue 40S ribosomes from degradation in homeostasis (*58,83*). Our findings suggest a working model wherein eIF2A interacts with G3BP1-USP10 complexes to counteract the deubiquitination and rescue of 40S subunits from otherwise programmed degradation.

Several lines of experimental evidence support a direct functional link between eIF2A and G3BP1-USP10 complexes in RQC. First, G3BP1 and USP10 consistently emerged as key interactors of eIF2A throughout our TurboID experiments. Second, zooming into the interaction with the 40S subunit, eIF2A interacted with the 40S proteins near the mRNA entry channel, including proximity to RPS2 and RPS3. Of note, eIF2A is a structural homolog of the eukaryotic initiation factor eIF3B (Fig. S5C), raising the possibility that eIF2A could replace eIF3B once it dissociates from 80S following translation initiation and early elongation (*79*). Third, loss of eIF2A resulted in reduced RPS3 ubiquitination upon ribosome stalling in both SCC and HEK 293T cells. Fourth, USP10 was required for the reduction in RPS3 monoubiquitination in *eIF2A knockouts* during stalling. Finally, loss of eIF2A resulted in markedly reduced 40S subunit turnover under stress in both SCC and HEK 293T cells. Collectively, our data support a model wherein eIF2A binds G3BP1-USP10 complexes to regulate RPS2 and RPS3 deubiquitination in response to ribosome stalling, effectively shifting the balance of the RQC pathway toward 40S subunit degradation. Our model is consistent with recent eCLIP data (*38*), which revealed that eIF2A predominantly binds coding sequences rather than 5’UTRs, in line with a role in RQC and subunit turnover rather than canonical initiation.

We can currently only speculate about the functional advantage of such a system. However, the capacity to regulate 40S deubiquitination and rescue may confer multiple potential benefits under specific cellular conditions. In homeostasis, the excess of USP10 relative to RNF10 ensures that deubiquitination is favored over degradation (*87*), which may avoid energetically costly subunit degradation. Although ribosome repair remains poorly studied, USP10 inhibition by eIF2A may provide cells with additional time to complete repair processes on RNA and ribosomes, particularly under cellular stress conditions. In this scenario, inhibition of eIF2A may result in the translation of aberrant mRNAs and the accumulation of faulty 40S ribosomes. In the context of iRQC, eIF2A may also act to delay re-initiation, potentially providing time for ternary complex acquisition, which is a process that could be particularly relevant under stress conditions marked by elevated levels of phosphorylated eIF2α. In this setting, as stalled preinitiation complexes are thought to eventually transition into elongating ribosomes (*83*), loss of eIF2A may lead to premature or aberrant re-initiation events.

Recent ribosome profiling experiments in yeast and human HeLa cells indicated that eIF2A has minimal impact on global translation, with only a few transcripts showing differential translation in homeostasis or following stress (*50,51*). Consistently, despite eIF2A’s critical role in early stages of tumor formation, its loss had minimal effects on translation in SCC cells in homeostasis *in vitro* (*46*). Our current selective 40S profiling data reinforce the notion that eIF2A is not a major player in translation initiation (Fig. 4). Given the identified link to 40S ribosome rescue in the context of RQC, it is perhaps not surprising that eIF2A depletion does not broadly alter the translational landscapes, as alterations in 40S turnover may not preferentially affect specific transcripts but rather contribute to global ribosome homeostasis. These downstream effects may only become apparent under conditions of chronic or low-level stress as opposed to short-term stress. Of note, pronounced eIF2A-dependent gene expression changes were observed in longterm pulsed SILAC under mild stress conditions, revealing over 350 differentially synthesized proteins (*46*).

Interestingly, elevated levels of both RNF10 and eIF2A are associated with shorter overall survival in patients with head and neck squamous cell carcinoma (*46*). Tumor samples also showed enhanced ubiquitination of RPS2 and RPS3, with a particularly strong increase at lysine 214, which is RPS3’s primary monoubiquitination site following ribosome stalling. These findings suggest heightened ribosome stalling or collision events in cancer and raise the possibility that RNF10-mediated 40S flagging, coupled with eIF2A-dependent inhibition of rapid 40S rescue, may provide a selective advantage to tumor cells. Future studies are needed to elucidate the role of 40S subunit turnover in cancer cells and to determine whether a heightened RQC response is particularly important for tumor progression.

Collectively, our study identifies an unexpected link between the alternative initiation factor eIF2A and G3BP1-USP10 complexes within the ribosome-associated quality control pathway, suggesting that eIF2A plays a major role in regulating 40S turnover in stress and warrants further investigations into how 40S turnover impacts tumorigenesis.

## Acknowledgements

We thank Aitor Garzia, Cindy Meyer and the Tuschl lab for discussions and sharing reagents, Tess Branon and the Ting lab for discussing the TurboID experimental design, Sendoel lab members for critical input on the manuscript, Witold Wolski, Antje Dittmann, Tobias Kockmann and the FGCZ for proteomics sample processing and data analysis, Catharine Aquino and the FGCZ for sequencing. The project was supported by the National Center of Competence in Research (NCCR) on RNA and Disease funded by the SNSF (grant number 205601), the European Research Council (ERC) under the European Union′s Horizon 2020 research and innovation programme (grant agreement No 759006), the SNSF Professorship grant (grant number 176825) and the UZH Candoc grant (grant number FK-22-045).

## Author contributions

M.Y. and A.S. conceived the project and designed the experiments. M.Y. conducted mass spectrometry experiments and collected data. M.Y., R.W. and K.N. performed 40S ribosome profiling. M.Y. and R.W. performed polysome gradients and luciferase assays. M.Y. and A.D. carried out co-immunopurification. M.Y. performed confocal microscopy imaging. F.V-F., D.T. and K.H. assisted with western blots and lentiviral preparations. K.H. assisted with cloning. M.O., C.D., H.Y. P.F.R., R.W. and K.H. assisted with generating cell lines and cell culture. M.Y., M.J., R.W., U.G. and A.S. conducted data analysis and interpretation. M.Y. and A.S. wrote the manuscript. A.S. supervised the project. All co-authors provided input on the manuscript.

## Data availability

The ribosome profiling data and the mass spectrometry proteomics data will be available upon publication.

## Materials and Methods

### Cloning TurboID gene constructs

V5::TurboID::NES_pCDNA3 mammalian expression plasmid (Addgene: #107169) and C1(1-29)::TurboID::V5_pLX304 (Addgene: #107175) ER targeting lentiviral plasmids were obtained from the Ting lab (*52*). C1(1-29)::TurboID::V5 was removed from the pLX304 lentiviral backbone and replaced with V5::TurboID::NES to create a cytosolic expression of TurboID in the V5::TurboID::NES-pLX304 backbone. CMV promoter was replaced with EF1-a promoter. eIF2A and tGFP was obtained from Eif2a (NM_001005509) Mouse tGFP-Tagged ORF Clone (Origene: #MG209105) and subsequently cloned into V5::TurboID::NES_pLX304 vector with a linker to generate to generate eIF2A::V5::TurboID::NES (C-term), V5::TurboID::NES::eIF2A (N-term) and tGFP::V5::TurboID::NES lentiviral constructs.

### Generating TurboID-expressing SCC cells by low-titer lentivirus transduction

Vesicular stomatitis virus (VSV-G) pseudotyped lentivirus was produced by calcium phosphate transfection of Lenti-X 293T (TaKaRa Clontech, #632180) cells with the TurboID constructs and the helper plasmids pMD2.G and psPAX2 (Addgene plasmids #12259 and #12260), following established procedures in the lab. After 46 hours, viral supernatants were harvested and filtered through a 0.45 µm pore-size filter (Sarstedt AG, #83.1826) to eliminate residual cells and debris. For lentiviral infections, SCC cells were seeded at a density of 1.5-2.5 × 10^5^ cells per well in a 6-well plate (Thermo Scientific Nunclon TM Delta Surface; #140675). 24 hours after, cells were infected with 300-600 µl of low-titer lentiviral supernatant supplemented with an infection mixture consisting of a 1:10 dilution of polybrene (10 mg/ml Sigma; #107689-100MG in PBS) in FBS [-], followed by centrifugation at 1100 *g* for 30 min at 37 °C in a Thermo Heraeus Megafuge 40R centrifuge. Infected cells were selected with Blasticidine (5 µg/ml) for 5 days.

### Primary keratinocyte isolation and culture

SCC cell lines were cultured in E-media with 0.05 mM Ca^2+^ and 15% Fetal bovine serum or fetal calf serum. Mouse squamous cell carcinoma (SCC) cells driven by HRas^G12V^; *Tgfbr2*^*fl/fl*^ were previously generated (*53*). *Eif2a* knockout *HRas*^*G12V*^; *Tgfbr2*^*-/-*^ SCC cells (clone 5A) and control sgRNA *HRas*^*G12V*^; *Tgfbr2*^*-/-*^ SCC (clone 4C) were generated previously (*46*). HEK 293T cells were cultured in standard Dulbecco’s Modified Eagle’s Medium (DMEM) supplemented with 10% FCS/FBS and 1% Penicillin/Streptavidin. All cell lines were maintained under standard culturing conditions at 37°C and 5% CO_2_.

### eIF2A knockout HEK 293T cell line

sgRNAs targeting eIF2A were designed using the CHOPCHOP (http://chopchop.cbu.uib.no) online tool and cloned into the lentiGuide-Puro plasmid. lentiGuide-Puro was a gift from Feng Zhang (Addgene plasmid #52963). The following guide sequences for eIF2A were used: sgRNA1: CTATAGTAGGCTGCATTACG; sgRNA2: GACTGAGAGACAG-TGTTTCG.

To obtain *eIF2A knockout* HEK 293T cells, parental cells were plated in 6 well plates. Notably, the parental HEK 293T cell line used in this study stably expresses an sfCherry3C(1-10)-P2A-Neomycin and mNG2(1-10)-P2A-EBFP construct which is not relevant for any conclusions of this study. Cells were transfected with plasmids encoding sgRNAs targeting eIF2A and a puromycin selection cassette in combination with a doxycycline inducible Cas9 construct using Lipofectamine 2000. After 24 hours, cells were selected with puromycin (3 µg/ml) for sgRNA and Cas9 expression induced with Doxycycline (4 µM) for 48 hours. Serial dilutions in 96-well plates were used to isolate single-cell clones. The loss of eIF2A in clonal cell lines was verified by western blotting.

### eIF2A KO rescue cells

V5::SBP::eIF2A construct was cloned into a pLKO-Blast backbone. Low-titer lentivirus production was conducted as mentioned above. *eIF2A knock-out* HEK 293T cells were infected and subsequently selected with Blasticidin (5 µg/ml) for a total of seven days. The partial rescue of V5::SBP::eIF2A expression in *eIF2A knockout* cells was verified by western blotting.

### TurboID-proximity labeling of SCC cells

TurboID proximity labeling was performed in *Hras*^*G12V*^; *Tgfbr2*^*-/-*^; *eIF2A*^*-/-*^ SCC cells expressing TurboID constructs. Cells were seeded at a density of 7 × 10^6^ cells/ml into 500 cm cell culture plates. To inhibit the activity of canonical eukaryotic translation initiation factor eIF2, cells were stressed with 50 µM sodium arsenite (0.05 M, Chemlab, #11963203) for 30 min followed by 1 hour co-incubation with 500 µM freshly prepared small molecule biotin (Sigma, #B4639-1G) in DMSO. Unstressed cells were used in parallel as a control and incubated with biotin for 1 hour. Biotinylation was stopped by washing three times with ice-cold 1x PBS (Gibco #10010-015). Cells were scraped off with 3 ml ice-cold 1x PBS and centrifuged for 5 min at 1500 rpm and 4°C. The cell pellet was lysed with ice-cold RIPA lysis buffer (Sigma #R0278) containing ice-cold RIPA lysis buffer (Sigma R0278) containing 1x Protease Inhibitor Cocktail (50x Promega, #G6521) for 10 min on ice and subsequently centrifuged at for 15 min at 4°C. Protein concentration was quantified using Pierce BCA Protein Assay Kit (Thermo Scientific #23225) according to manufacturer guidelines. 10% of the cell lysate was stored at –80°C for further western blot experiments. Fresh protein lysate was subjected to further capture of eIF2A interactors.

### Capturing eIF2A interactors with streptavidin-coated magnetic beads

The Pierce Streptavidin-coated Magnetic Beads (Thermo Scientific, #88817) were mixed thoroughly by pipetting up and down, avoiding vortexing. Per 3 mg of protein sample, 225 µl of streptavidin-coated magnetic beads were transferred into a 1.5 ml Protein LoBind Eppendorf tube and placed on a magnetic stand. The supernatant was carefully aspirated, the protein LoBind tube was removed from the magnetic stand, 1 ml of ice-cold RIPA lysis buffer (Sigma #R0278) containing 1x Protease cocktail inhibitor (50x Promega, #G6521) was added onto beads and the contents were mixed by pipetting. Following a 60-second incubation, the tube was returned to the magnetic stand, the supernatant was aspirated and the washing step was repeated. After the second wash, the beads were resuspended in 225 µl of ice-cold RIPA lysis buffer (Sigma #R0278) containing 1x Protease cocktail inhibitor (50x Promega, #G6521).

Protein lysates (3 mg) were added to magnetic beads. The total volume of bead-slurry was brought up to 1.5 ml with RIPA buffer, including the bead volume. The mixture was incubated on a rotating shaker (Stuart EW-07650) for 1 hour at room temperature at speed 4, followed by overnight incubation in a cold room at speed 10.

After incubation, the tubes were placed on a magnetic stand, and the flow-through was collected in new protein LoBind Eppendorf tubes for further western blot analysis. Washing steps were then carried out using protein LoBind pipette tips, with each wash incubated for 60 seconds before magnetic separation and removal of the supernatant. The beads were washed twice with 1 ml of ice-cold RIPA lysis buffer (Sigma #R0278) containing 1x Protease cocktail inhibitor (50x Promega, #G6521), once with 1 ml of 1 M KCl, once with 1 ml of 0.1 M Na_2_CO_3_, and once with 1 ml of 2 M urea in 10 mM Tris-HCl (pH 8.0). This was followed by two additional washes with ice-cold RIPA lysis buffer (Sigma #R0278) containing 1x Protease Inhibitor Cocktail (50x Promega, #G6521) and a second wash with 2 M urea in Tris-HCl. The beads were then washed three times with PBS, transferring to new protein LoBind tubes at each step.

Finally, the beads were resuspended in 1 ml of 50 mM Tris and transferred into a new protein LoBind Eppendorf tubes. A 2.5% aliquot of the bead slurry (25 µl) was removed for western blotting. The remaining slurry was stored at +4 °C for mass spectrometry.

### 40S TurboID-proximity labeling of SCC cells

40S TurboID proximity labeling was performed as described above, with an additional step of sucrose density gradient fractionation after cell lysis before streptavidin-coated magnetic bead purifications. Briefly, 5.5 × 10^6^ cells were seeded in 500 cm culture plates (2 x per condition). After 30 minutes of biotinylation and harvesting with ice-cold RIPA lysis buffer (Sigma #R0278) containing 1x Protease Inhibitor Cocktail (50x Promega, #G6521), cell lysates from unstressed and sodium arsenite-stressed samples were layered onto a 10–50% sucrose gradient made in mammalian polysome gradient buffer (20 mM Tris-HCl pH 7.4; 150 mM NaCl; 5 mM MgCl_2_; 100 µg/ml cycloheximide) and subjected to ultracentrifugation at 41,000 rpm for 2 hours. Polysome fractions were then collected using the Biocomp Density Gradient Fractionation System. Then, collected fractions were pooled together into 2 ml protein LoBind Eppendorf tubes for 40S and 15 ml protein LoBind falcons for 80S + polysome fractions: Fraction 5 – 40S ribosome; Fraction 8 to 15 – 80S ribosome + polysomes.

Subsequently, streptavidin-coated magnetic beads were prepared as described above and 25 µl of beads and 50 µl of beads were added into 40S and 80S samples, respectively. The tubes were subsequently filled up to 2 ml and 15 ml with mammalian polysome buffer (20 mM Tris-HCl pH 7.4; 150 mM NaCl; 5 mM MgCl_2_; 100 µg/ml cycloheximide), including the bead volume. The mixture was incubated on a rotating shaker (Stuart EW-07650) for 1 hour at room temperature at speed 4, followed by overnight incubation in a cold room at speed 10. After incubation, the procedure was continued with salt wash steps as described above and followed by LC-MS/MS preparation.

### LC-MS/MS sample processing and on-beads digestion

Mass spectrometry analysis was carried out at Functional Genomics Center Zurich (FGCZ). Samples were prepared using an on-beads digestion protocol. The beads were first washed twice with 100 µl of digestion buffer (10 mM Tris, 2 mM CaCl_2_ pH 8.2), and the wash solutions were discarded after each step. Subsequently, 45 µl of the same digestion buffer was added to the beads. Reduction and alkylation were carried out using TCEP and chloroacetamide under standard conditions. Following this, 5 µl of trypsin (100 ng/µL in 10 mM HCl) was added to each sample. When necessary, the pH was adjusted to 8.0 to optimize enzyme activity. Digestion was performed overnight at 37 °C.

After incubation, the supernatants were collected and peptides were extracted from beads using 150 µl of 0.1% Trifluoroacetic acid (TFA) in 50% Acetonitrile. The supernatants were combined and subsequently dried by vacuum centrifugation in preparation for downstream analysis.

### Data acquisition

The digested peptide samples were dried and subsequently reconstituted in 20 µl of ddH_2_O containing 0.1% formic acid. The resuspended samples were transferred into autosampler vials for liquid chromatography-tandem mass spectrometry (LC-MS/MS) analysis. An injection volume of 1 µl was loaded onto an M-class UPLC system (Waters) coupled to a Fusion Lumos mass spectrometer (Thermo Scientific) for peptide separation and detection.

### LC-MS/MS data analysis

The LC-MS/MS data were processed using the MaxQuant and FragPipe proteomics pipeline (https://fragpipe.nesvilab.org/). To assess potential interactions between observed proteins (potential prays) and the bait protein, the resulting data were further analyzed using SAINTexpress: Significance Analysis of INTeractome – Express (http://saint-apms.source-forge.net/Main.html). The Bait-Prey interaction candidate lists were filtered using a Bayesian false discovery rate (BFDR) and SAINT score.

### Sucrose density gradient fractionation

Cells were seeded into 15 cm plates at a density of 70-90% confluency on the day of collection. Before harvesting, cells were incubated with different stressors (50 µM sodium arsenite, 1 µM anisomycin, and 100 µM anisomycin) for the duration indicated in each polysome gradient fractionation. Control cells were seeded in parallel and replaced with fresh media for the same amount of time. For the isolation of polysome fractions, cell lysates were prepared using a lysis buffer containing 20 mM Tris-HCl pH = 7.4, 150 mM NaCl, 5 mM MgCl_2_, 1% Triton X-100, 0.5% NP40, 1 mM DTT, and 100 µg/ml cycloheximide. The lysates were layered onto a 10–50% sucrose gradient made in mammalian polysome gradient buffer (20 mM Tris-HCl pH 7.4; 150 mM NaCl; 5 mM MgCl_2_; 100 µg/ml cycloheximide) and subjected to ultracentrifugation at 41,000 rpm for 2 hours. Polysome fractions were then collected using the Biocomp Density Gradient Fractionation System and mixed with a 2X protein sample (100 mM Tris-HCl pH 6.8, 4% SDS, 20% glycerol, 0.2 M DTT) for further western blot analysis.

### Western blot

Cells were seeded at a density of 70-90% confluency. On the day of the experiment, cells were incubated with different stressors (50 µM sodium arsenite for 2 hours, 1 µM anisomycin for 30 minutes, 100 µM anisomycin for 30 minutes, 2 µM MG132 for 6 hours, 10 µM spautin for 10 hours, 1 µM SAR405 for 10 hours, 3.2 µM rocaglamide for 2 hours and 2 µg/ml harringtonine for 1-2 hours) according to the experimental setup. Control cells were prepared in parallel by replacing them with fresh media.

Following incubation, cells were washed with ice-cold 1x PBS (Gibco, #10010-015), collected by scraping, and lyzed in RIPA buffer supplemented with 1x Protease Inhibitor Cocktail (50x, Promega, G6521) for 10 minutes on ice. Protein concentrations were measured using the Pierce BCA Protein Assay Kit (Thermo Scientific, #23225), following the manufacturer’s instructions. For sample preparation, protein lysates were mixed with 1x NuPAGE LDS Sample Buffer (4x, Invitrogen, #NP0007) and 1x NuPAGE Sample Reducing Agent (10x, Invitrogen, #NP0009), then denatured by heating at 90 °C for 10 minutes. Samples were loaded on a 4–12% NuPAGE Bis-Tris Gel (12-well, Invitrogen, #7001691) using 1x NuPAGE MOPS SDS Running Buffer (20x, Invitrogen, #NP0001).

Subsequently, proteins were transferred to a nitrocellulose membrane (Cytiva Amersham Protran 0.45 µm NC, #10600002) by tank transfer at 4°C, 30V for 90 minutes. Membranes were blocked in 5% BSA for 1 hour or 5% skim milk for 2 hours at room temperature. Primary antibodies were incubated overnight at 4°C, followed by 3 washes with TBS-Tween 0.1%. Secondary antibodies were incubated for 2 hours at 4 °C followed by 3 washes with TBS-Tween 0.1% before developing with freshly mixed ECL solutions (Amersham Cytiva, RPN2209 or Thermo Scientific #34094). Western blot bands were quantified using ImageJ version 2.1.0.

### Antibodies

The following antibodies were used: Anti-EIF2A/CDA02 polyclonal antibody (Proteintech, #11233-1-AP), eIF2A monoclonal antibody (1G7H4) (Proteintech, #66482-1-IG), G3BP1 polyclonal antibody (Proteintech, #13057-2-AP), Purified mouse anti-human G3BP1 monoclonal antibody (BD Biosciences, #611126), SERBP1 monoclonal antibody (M02), clone 1G5-2D7 (Abnova, #H00026135-M02), Anti-USP10 antibody (Abcam, #ab70895), RPS2 polyclonal antibody (Bethyl, #A303-794A-T), RPS3 polyclonal antibody (Bethyl, #A303-840A), RPS10 antibody (Abcam, #ab151550), ATF-4 (D4B8) rabbit mAb (Cell Signalling, #11815), eIF2S1 polyclonal antibody (Proteintech, #11170-1-AP), V5 Tag monoclonal antibody (Invitrogen, #R960-25), Streptavidin, horseradish peroxidase conjugate (Invitrogen, #S911), β-Actin monoclonal antibody (8H10D10) (Thermo Scientific, #MA5-15452), GAPDH monoclonal antibody (1E6D9) (Proteintech, #60004-1-Ig), Vinculin polyclonal antibody (Proteintech, #26520-1-AP), Tubulin antibody (Cell Signalling, #2144S), Anti-rabbit IgG HRP-linked antibody (Cell Signaling, #7074) and Anti-mouse IgG HRP-linked antibody (Cell Signaling, #7076), Goat anti-Mouse IgG (H+L) Highly Cross-Adsorbed Secondary Antibody, Alexa Fluor 594 (Invitrogen, #A-11032), Goat anti-Rabbit IgG (H+L) Cross-Adsorbed Secondary Antibody (Alexa Fluor 488) (Invitrogen, #A-11008).

### Immunopurification

Cells were seeded onto 15 cm culture plates at a density of 70-90% on the day of immunopurification. Cells were washed twice with ice-cold 1x PBS, scraped off and collected into a 15 ml protein LoBind falcon. After centrifugation for 5 minutes at 4°C and 1500 rpm, the cell pellet was lyzed with a mild lysis buffer containing 150 mM NaCl, 50 mM Tris pH 7.5, 0.05% NP-40, 5% glycerol, 1x Protease Inhibitor Cocktail. After incubation on ice for 10 min and centrifugation (14,000 x g, 10 min, 4 °C), protein concentration was assessed by a BCA assay. For each immunoprecipitation (IP), 650 µg of total protein (input) was incubated with 65 µl of Dynabeads Protein G (Thermo Scientific, #10004D) (pre-washed 3x in mild lysis buffer) and 2.5 µg of eIF2A polyclonal antibody, rotating overnight at 4 °C. Beads were washed twice with wash buffer containing 0.05% NP-40 (150 mM NaCl, 50 mM Tris pH 7.5, 5% glycerol, 0.05% NP-40), then twice with NP-40–free wash buffer (150 mM NaCl, 50 mM Tris pH 7.5, 5% glycerol). Bead-lysate slurry was boiled in a 2X protein sample buffer (100 mM Tris-HCl pH 6.8, 4% SDS, 20% glycerol, 0.2 M DTT) at 95 °C for 5 minutes, followed by centrifugation (10,000 x g, 3 minutes, 4 °C). The supernatant protein (IP) was further subjected to western blot analysis.

For western blotting, 5 µg of input was prepared to a final volume of 20 µl and heated in a 2X protein sample buffer (100 mM Tris-HCl pH 6.8, 4% SDS, 20% glycerol, 0.2 M DTT) to 95°C for 5 min. For the eIF2A immunoprecipitation western blot, 20 µl of input and 25 µl of IP were loaded onto 4-12% NuPAGE 12-well Bis-Tris Gel (Invitrogen, #7001691). For co-immunopurification of the interactors, 4 µl of input and 25 µl of IP were loaded onto 4-12% NuPAGE 12-well Bis-Tris Gel (Invitrogen, #7001691).

### Immunofluorescence

SCC and HaCat cells were seeded onto coverslips in a 24-well plate. HEK 293T cells were seeded onto poly-L-lysine-coated coverslips in a 24-well plate. The coating was performed using a 1:10 dilution of poly-L-lysine in ddH_2_O for 1 hour at room temperature, followed by three 1x PBS washes. Cells were cultured for 24-48 h before processing. For stress conditions, cells were incubated with sodium arsenite (500 µM for HEK 293T and HaCaT cells; 50 µM for SCC cells). Cells were fixed with 4% paraformaldehyde (16% formaldehyde solution (w/v), methanol-Free, Thermo Scientific #28908) in 1x PBS for 15 min at room temperature and washed three times in 1x PBS. Permeabilization was carried out using 0.3% Triton X-100 in 1x PBS for 15– 20 min. Blocking was performed using a buffer containing 1% BSA, 1% fish skin gelatin, 2.5% horse serum, and 0.3% Triton X-100 in 1x PBS for 1 hour at room temperature. Primary antibodies were diluted in antibody dilution buffer (0.3% Triton X-100 and 1% BSA in 1x PBS) and incubated overnight at 4 °C. After three times 5-minutes washes in PBS-T (0.3% Triton X-100), coverslips were incubated with secondary antibodies conjugated with fluorophores for 45 min at room temperature in the dark. Nuclei were stained using DAPI (0.02 ng/µl) for 15 minutes, followed by two additional PBS-T washes. Coverslips were mounted with homemade Antifade mountant, airdried under tissue paper for 2–3 hours, sealed with nail polish, and stored at 4 °C protected from light. Imaging was performed with 63x magnification on a Leica SP8 confocal microscope (Leica, Germany).

### Anisomycin-induced ribosome collision assay

SCC and HEK 293T BB cells were seeded in 15 cm culture plates and incubated with either intermediate dose (1 µM) or high dose (100 µM) Anisomycin for 30 minutes. In parallel, cells were incubated with 50 µM sodium arsenite for 2 hours. Control cells were kept untreated. After incubation, cells were washed with ice-cold PBS, scraped off and lyzed with polysome lysis buffer (20 mM Tris-HCl pH = 7.4, 150 mM NaCl, 5 mM MgCl_2_, 1% Triton X-100, 0.5% NP40, 1 mM DTT, and 100 µg/ml cycloheximide) for 10 minutes on ice. After centrifugation for 10 minutes at 4°C, the protein concentrations were quantified and RPS3 ubiquitination levels were measured by western blotting.

### Collision-inducing luciferase reporter assay

HEK 293T BB cells were seeded in a 6-well plate and transiently transfected with a collision-inducing *Renilla* luciferase-16-Lysine (corresponding to polyA) reporter or a control *Renilla* luciferase-stop codon-16-Lysine reporter. *Firefly* luciferase was co-transfected as an internal control (3:1 ratio of Rluc:Fluc). The luminescence signal was measured using Dual-Luciferase Reporter Assay (Promega, #E1910). Briefly, after 24 hours of transient transfection, cells were washed with 1x PBS, scraped off and lysed with 1x passive lysis buffer (PLB) in distilled water. After centrifugation, 5 µl of supernatant cell lysate was transferred into clear bottom, white 96-well plates. Then, 25 µl LAR II (Luciferase Assay Substrate in Luciferase Assay Buffer II) reagent was added to cell lysate. *Renilla* luciferase activity was measured in a TECAN reader. Subsequently, 50 µl 1x Stop & Glo reagent (50x Stop & Glo Substrate in Stop & Glo Buffer) was added and the firefly luciferase activity was measured. Three biological replicates were conducted with three technical replicates each.

### RNF10 overexpression

Human FLAG-HA tagged wild-type and mutant RNF10 (FLAG::HA::RNF10^Wildtype^, FLAG::HA::RNF10^C228S^, FLAG::HA::RNF10^C225S^ FLAG::HA::RNF10^C240S/C245S^, FLAG::HA::RNF10^C225S/C228S^) overexpression constructs were generously provided by the Tuschl lab. HEK 293T BB cells were seeded onto a 6-well plate and transiently transfected with 2.5 µg of overexpression constructs using Lipofectamine 3000 Transfection Reagent (Thermo Scientific) according to manufacturer’s guidelines. Control cells were transiently transfected with H2B-GFP. After 48-96 hours of transfection, cells were lyzed with a polysome lysis buffer (20 mM Tris-HCl pH = 7.4, 150 mM NaCl, 5 mM MgCl_2_, 1% Triton X-100, 0.5% NP40, 1 mM DTT, and 100 µg/ml cycloheximide) for 10 minutes on ice. After centrifugation for 10 minutes at 4°C, the protein concentrations were quantified and RPS3 ubiquitination levels were measured by western blotting.

### CRISPR-mediated Usp10 knockout

Single guide RNAs (sgRNAs) targeting Usp10 were designed using the Broad Institute’s CRISPRko sgRNA Designer tool (broadinstitute.org). Four sgRNA targeting mouse Usp10 were cloned into a pLentiCRISPR-blasticidin (cloned from pLentiCRISPRv2-mCherry (Addgene #99154)) backbone: sgRNA1: AAGTCATCGAACCTAGTGAG, sgRNA2: GGAAA-GCAACTCTAACGCAG sgRNA3: GGCATGGCCGTTGACCAGGG, sgRNA4: TCGTGAGAGATATCCGCCCA.

Low-titer lentivirus was prepared and *HRas*^*G12V*^; *Tgfbr2*^*-/-*^ SCC cells were infected to generate *Usp10 knockout* and *Eif2a; Usp10 double knockout* lines. Cells were then selected with blasticidin (5 µg/ml) and clonal cell lines were isolated using fluorescence-activated cell sorting (FACS). The frameshift alleles were confirmed with a Tide assay for sg4. Loss of USP10 expression in knockout cells was confirmed by western blot analysis.

### Stable isotope labeling by amino acids in cell culture (SILAC)

Heavy and light isotope-containing cell culture mediums were prepared according to the manufacturer’s protocol (SILAC Protein Quantitation Kit (Trypsin) –RPMI 1640 Kit #A33973). Parental and *eIF2A knockout* HEK 293T cells were cultured in the light isotope-containing medium in a 10 cm dish at a density of 20-30% confluency. After 24 hours, cells were switched to heavy isotope-containing medium, and protein samples were collected at 6 hours, 12 hours, and 24 hours post-switch. HEK 293T cells were co-incubated with 5 µM low dose sodium arsenite during heavy isotope labeling. In parallel, unstressed cells were incubated only with a heavy isotope.

SCC cells were cultured in E-lo media, seeded in light isotope-containing media for 24 hours and co-incubated with 15 µM sodium arsenite during heavy isotope labeling. Protein samples were collected at 6, 12 and 24 hours after the media switch and prepared for proteomics analysis.

Briefly, cells were washed three times with ice-cold PBS, scraped off and collected into protein LoBind falcons. After pelleting, cells were lysed in 150 µl RIPA buffer containing 1X Protease Inhibitor Cocktail, incubated for 10 minutes on ice and centrifuged (14,000 x g, 10 min, 4°C). The protein supernatant was subjected to mass spectrometry at the FGCZ.

### SILAC proteomics sample preparation and data acquisition

Protein concentrations were measured using the Lunatic UV/Vis absorbance spectrophotometer (Unchained Labs). Proteins were reduced and alkylated by adding Tris(2-carboxyethyl)phosphine and 2-Chloroacetamide to a final concentration of 5 mM and 15 mM, respectively. The samples were incubated for 30 minutes at 30° C; 700 rpm and were light-protected. The samples were diluted with pure ethanol to reach a final concentration of 60% EtOH (v/v).

The following steps were carried out on a KingFisher Flex System (Thermo Scientific): adding the corresponding amount of carboxylated magnetic beads (hydrophobic and hydrophilic) to the samples. After binding the proteins to the beads for 30 minutes at room temperature, beads were washed three times with 80% EtOH.

For the enzymatic digestion, the beads were added to trypsin in 50 mM TEAB. Samples were digested overnight at 37 °C. The remaining peptides were extracted from beads with H_2_O. The two elutions were combined and dried down.

The digested samples were dissolved in aqueous 3% Acetonitrile with 0.1% formic acid, and the peptide concentration was estimated with the Lunatic UV/Vis absorbance spectrometer (Unchained Lab). Peptides were separated on an M-class UPLC and analyzed on an Iontrap mass spectrometer (LC-MS/MS) (Thermo Scientific).

### SILAC proteomics data analysis

The acquired MS data were processed using the Fragpipe v19 (MSFragger search engine, PMID: 28394336). Spectra were searched against Mus musculus protein database. Carbamidomethylation of cysteine was set as a fixed modification, while oxidation of methionine and protein N-terminal acetylation were set as variable modifications.

### Ribosome profiling

Cells were plated on 15 cm dishes 48 hours before harvesting. Cells were incubated for 2 hours with 50 µM sodium arsenite (stress) or left untreated (control) prior to harvesting.

40S Ribosome profiling (*78,79*): Cells were briefly washed with ice cold wash buffer (10 mM MgCl_2_, 800 µM cycloheximide in 1x PBS) and crosslinked with crosslinking solution (10 mM MgCl_2_, 200 µM cycloheximide, 0.025% formalydehyde, 0.5 mM DSP in 1x PBS) for 15 minutes at room temperature gently shaking. The reaction was stopped using quenching solution (10 mM MgCl_2_, 200 µM cycloheximide, 300 mM glycine in 1x PBS) for 5 min at room temperature gently shaking. All liquid was thoroughly removed and cells were lysed in 500 µl lysis buffer (20 mM Tris-HCl pH=7.4, 150 mM NaCl, 5 mM MgCl_2_, 1 mM DTT, 0.5% NP40, 35 µM cycloheximide, 1% TritonX-100). Lysates were incubated on ice for 5 minutes, spun down 5 minutes at 20,000 x g at 4 °C and supernatants were collected. 300 µl of the lysates were treated with 4 µl RNase I for 20 minutes at room temperature and subsequently run over a 10-50% sucrose gradient. Fractions corresponding to the 40S ribosomal subunit were collected and pooled to obtain an approx. volume of 500 µl sample. Samples were treated with 55 µl crosslink removal solution (10% SDS, 100 mM EDTA pH=8, 50 mM DTT) and 600 µl Acid-Phenol-Chloroform for 45 minutes at 65°C vigorously shaking. Afterwards, samples were cooled down on ice and spun down 5 minutes at 20,000 x g at 4°C. The upper aqueous phase was collected, mixed with 3 volumes of Trizol LS and 4 vol 100% ethanol and 40S footprints were isolated using the Direct-zol RNA Miniprep Kit.

80S Ribosome profiling: Cells were briefly washed with ice cold wash buffer (35 µM cycloheximide in 1x PBS), all liquid was thoroughly removed and samples flash frozen in liquid nitrogen. Cells were lysed in 500 µl lysis buffer (20 mM Tris-HCl pH=7.4, 150 mM NaCl, 5 mM MgCl_2_, 1 mM DTT, 0.5% NP40, 35 µM cycloheximide, 1% TritonX-100, 2 U/µl TurboDNase). Lysates were incubated on ice for 5 minutes, spun down 5 minutes at 20,000 x g at 4 °C and supernatants were collected. 150 µl of the lysates were treated with 2 µl RNase I for 45 minutes at room temperature. The reaction was stopped by adding 4 µl SUPERase Inhibitor and placing the samples on ice. 80S footprints were subsequently isolated using Illustra MicroSpin S-400 HR columns, mixed with 3 volumes of Trizol LS and 4 vol 100% ethanol and purified using the Direct-zol RNA Miniprep Kit.

40S and 80S Ribo-seq libraries were prepared largely following the protocol by McGlincy & Ingolia (*88*). Briefly, ribosome footprints were excised and extracted from a 15% TBE-Urea gel; 80S: size range 20-35 nts; 40S: size range 20-80 nts. Samples were subjected to an end-healing reaction for 1 hour at 37 °C and linkers ligated for 3 hours at 22 °C (for details see McGlincy & Ingolia (2017)). Ribosomal RNA was depleted using custom biotinylated rRNA depletion oligos and MyOne Streptavidin dynabeads. Reverse transcription was performed using SuperScript IV. cDNA was excised and extracted from a 15% TBE-Urea gel and circularized using CircLigase II for 1 hour at 60 °C. cDNA was amplified using Phusion High-Fidelity DNA polymerase using an individual index per sample and purified over an 8% TBE gel (for details see McGlincy & Ingolia (2017)).

**Figure S1:**
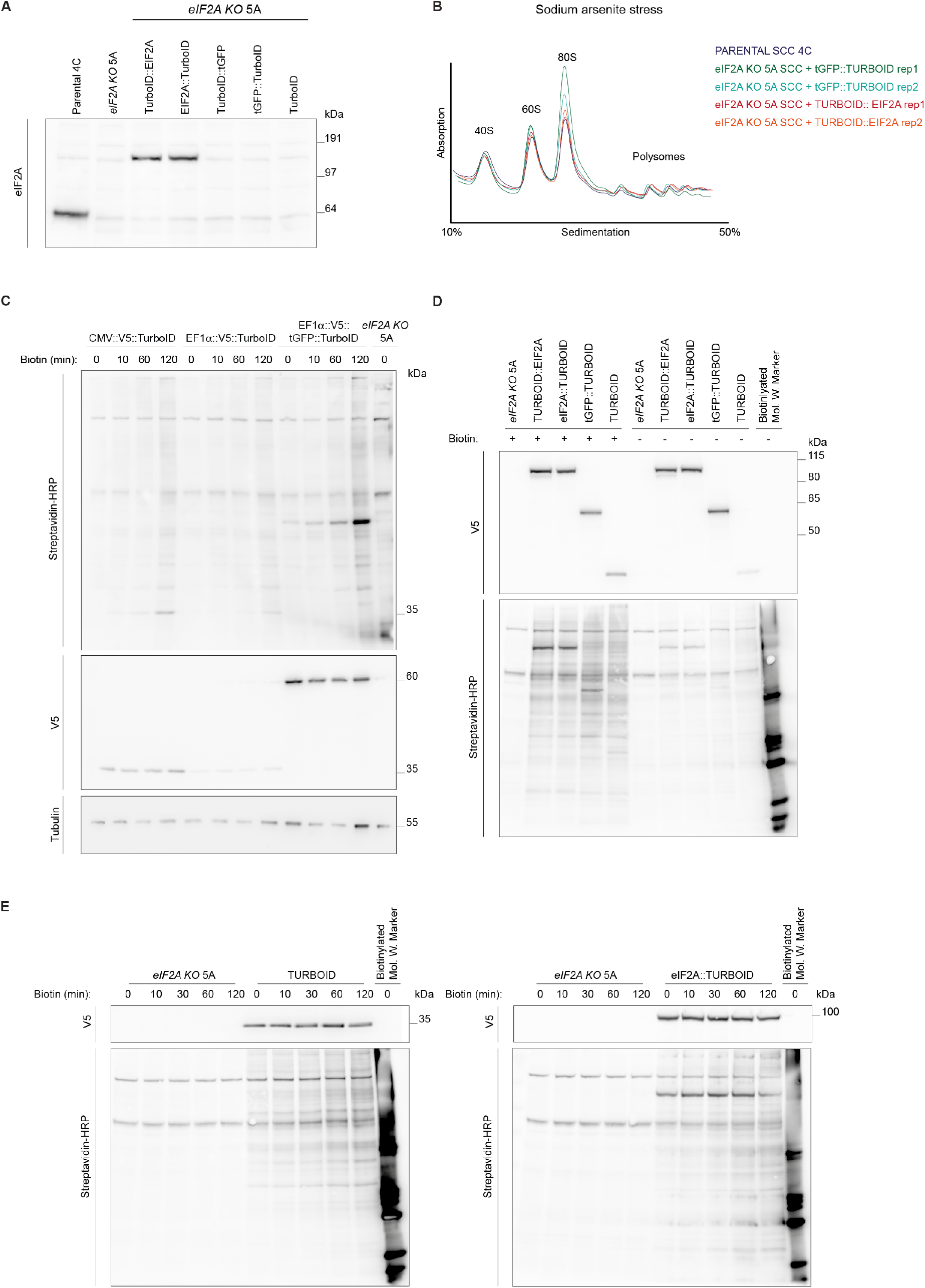
TurboID-based proximity biotinylation strategy in SCC cells. **(A)** Western blot displaying the expression of eIF2A TurboID constructs in SCC cells. eIF2A expression was restored in *eIF2A knockout* cells using lentiviral transduction of either eIF2A::TurboID or TurboID::eIF2A, with levels comparable to endogenous eIF2A. **(B)** Polysome gradient profiles of SCC cells expressing TurboID constructs. The expression of eIF2A::TurboID or TurboID::eIF2A in *eIF2A knockout* cells partly rescued the mild defects in subunit stoichiometry to levels comparable with parental cells. **(C)** Western blot analysis of biotinylation in SCC cells expressing TurboID constructs, using streptavidin-HRP antibodies. Cells were incubated with biotin for 10 to 120 minutes. Constructs driven by the EF1α promoter showed stronger biotinylation signals compared to those under the CMV promoter. All subsequent TurboID constructs were therefore expressed under the EF1α promoter. TurboID constructs were detected with V5 antibodies; Tubulin served as a loading control. **(D)** Western blot showing expression of TurboID constructs (V5) and biotinylation after 1 hour in SCC cells. A biotinylated molecular weight marker was used as a positive control. **(E)** Biotinylation time course in *eIF2A knockout* SCC cells expressing either TurboID alone (left panel) or eIF2A::TurboID (right panel). TurboID construct expression was detected with a V5 antibody and biotinylation was monitored using streptavidin-HRP.

**Figure S2:**
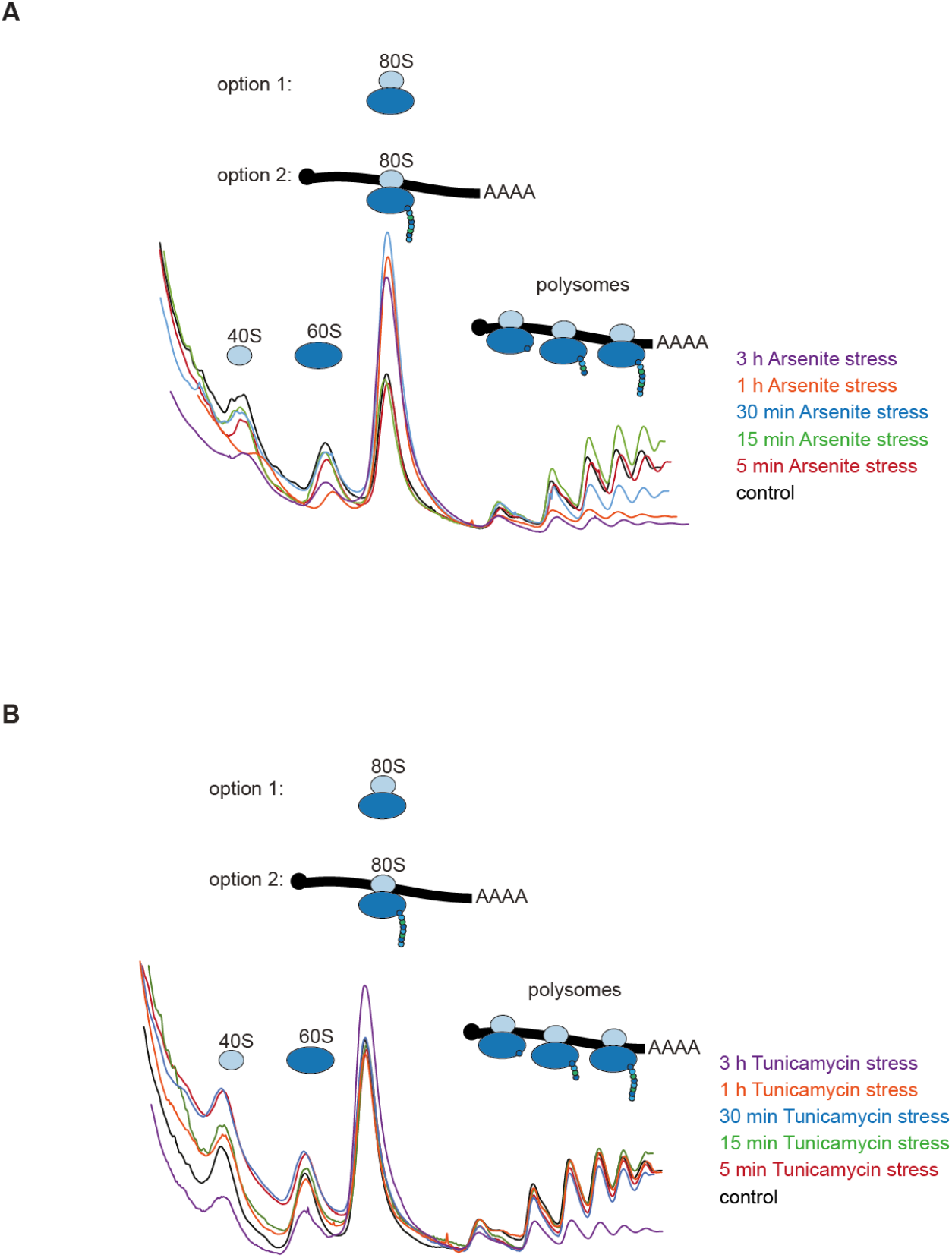
Polysome gradients upon stress. **(A)** Polysome gradient profiles of parental SCCs subjected to sodium arsenite stress over a time course from 5 minutes to 3 hours. **(B)** Polysome gradient profiles of parental SCCs treated with tunicamycin over a time course from 5 minutes to 3 hours.

**Figure S3:**
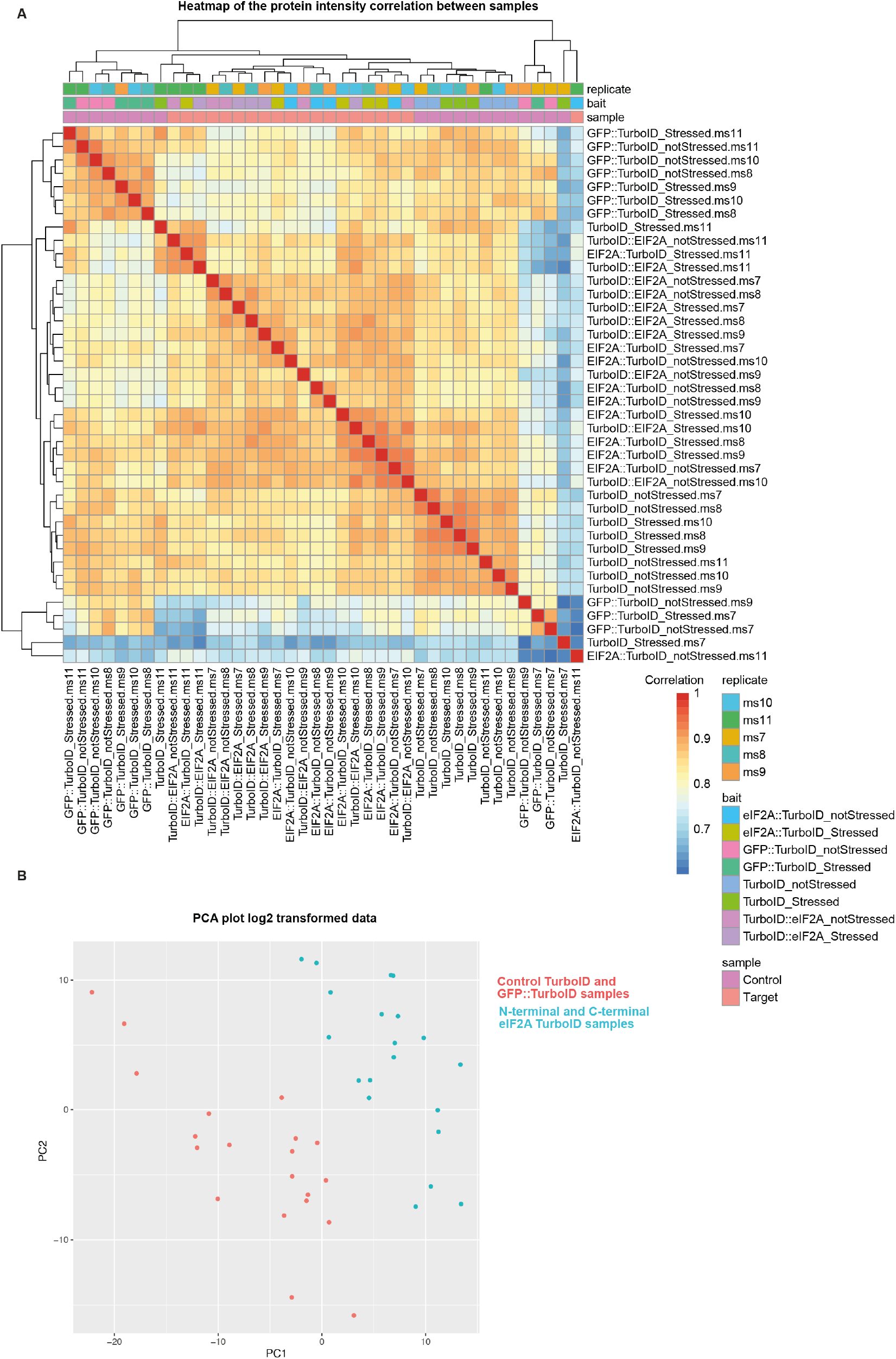
Heatmap of protein intensity correlation. **(A)** Heatmaps showing protein intensity correlations across five biological replicates from the main TurboID-based mapping of eIF2A interactors. **(B)** Principal component analysis (PCA) of all samples in the main TurboID proximity labeling. Control and eIF2A-expressing samples form distinct clusters.

**Figure S4:**
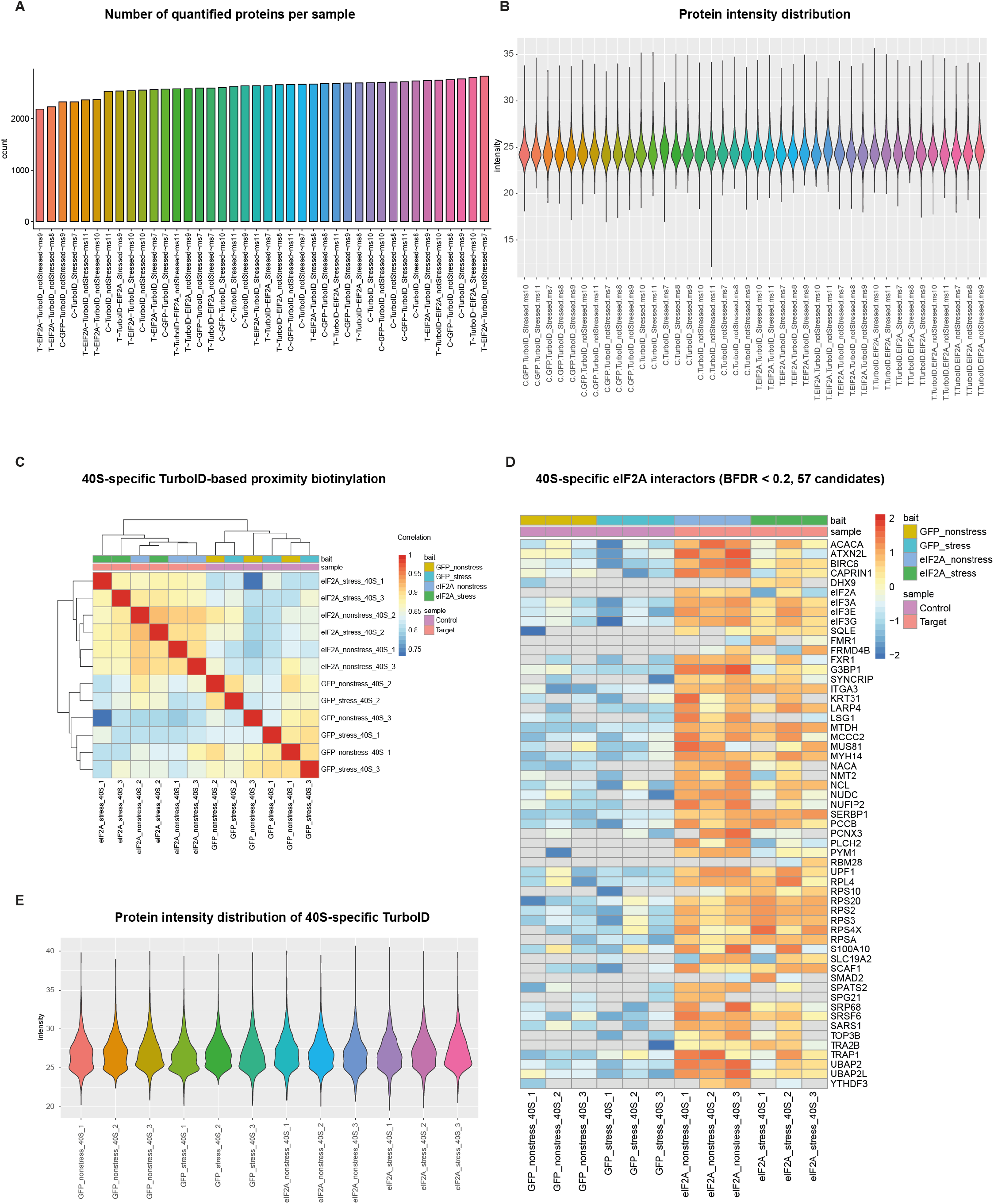
Scatterplot of samples. **(A)** Bar plots display the number of quantified proteins per sample across replicates in TurboID proximity labeling. **(B)** Violin plot showing the distribution of protein intensities. **(C)** Heatmap illustrating protein intensity correlations among 3 biological replicates from the 40S-specific TurboID-based interactome analysis of eIF2A. **(D)** Heatmap displays the intensity of top 40S-specific eIF2A interactors (57 interactors with BFDR < 0.2) across the 3 replicates. **(E)** Violin plot showing the distribution of protein intensities of the 40S-specific TurboID-based mapping of eIF2A interactors.

**Figure S5:**
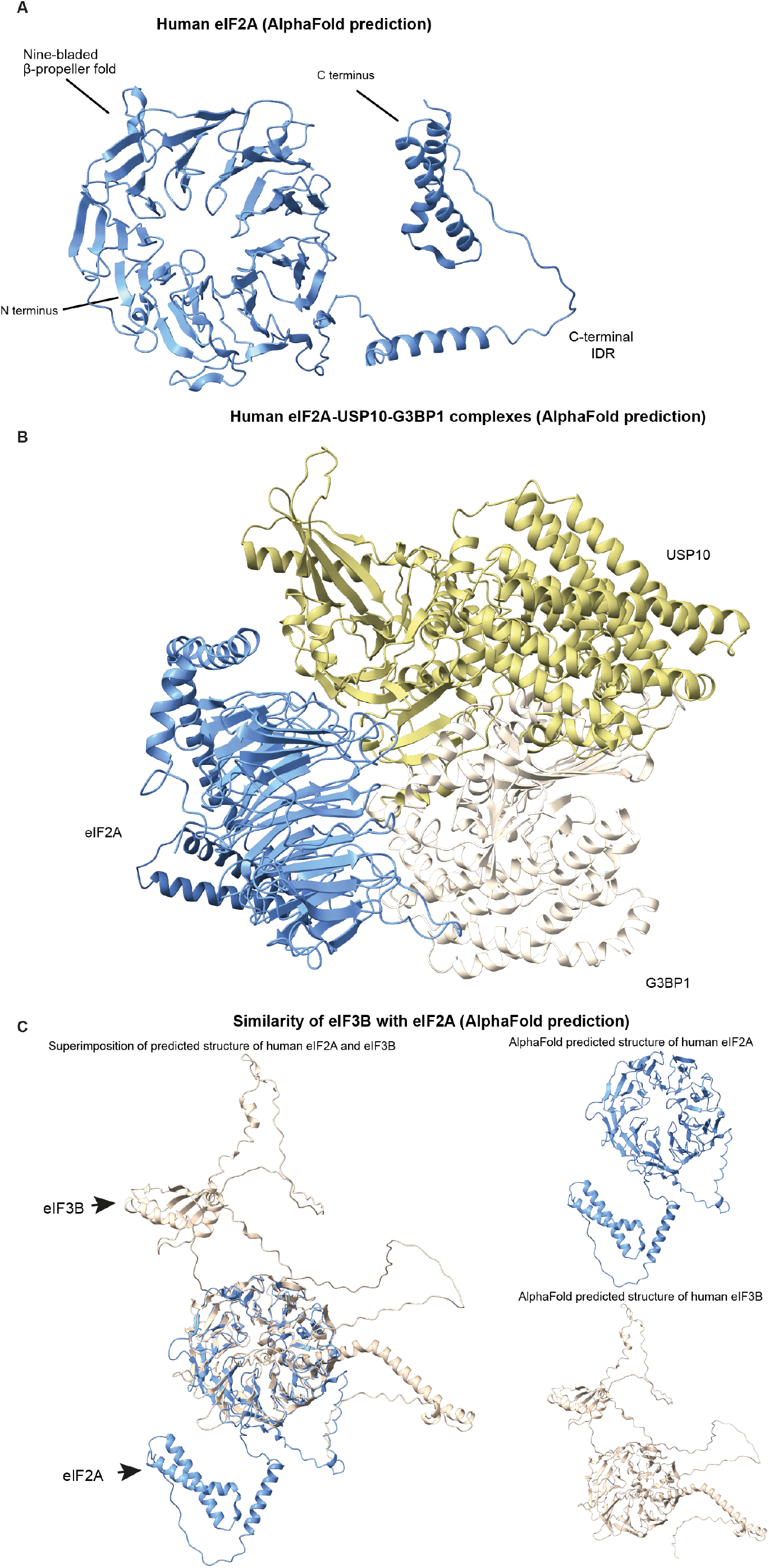
AlphaFold predictions. **(A)** The AlphaFold-predicted structure of the full-length human eIF2A. The N-terminus contains a nine-bladed β propeller, while the C-terminal part is more flexible and contains an intrinsically disordered region (IDR). **(B)** Predicted AlphaFold-multimer structure of human eIF2A-G3BP1-USP10 complex. ipTM (inter-protein Template Model) score = 0.25 and pTM (per-chain Template Model) score = 0.36, with ipTM indicating the likelihood of correct inter-protein interactions and pTM providing a per-chain accuracy estimate. **(C)** Superimposition of the predicted AlphaFold structure of human eIF2A and eIF3B.

**Figure S6:**
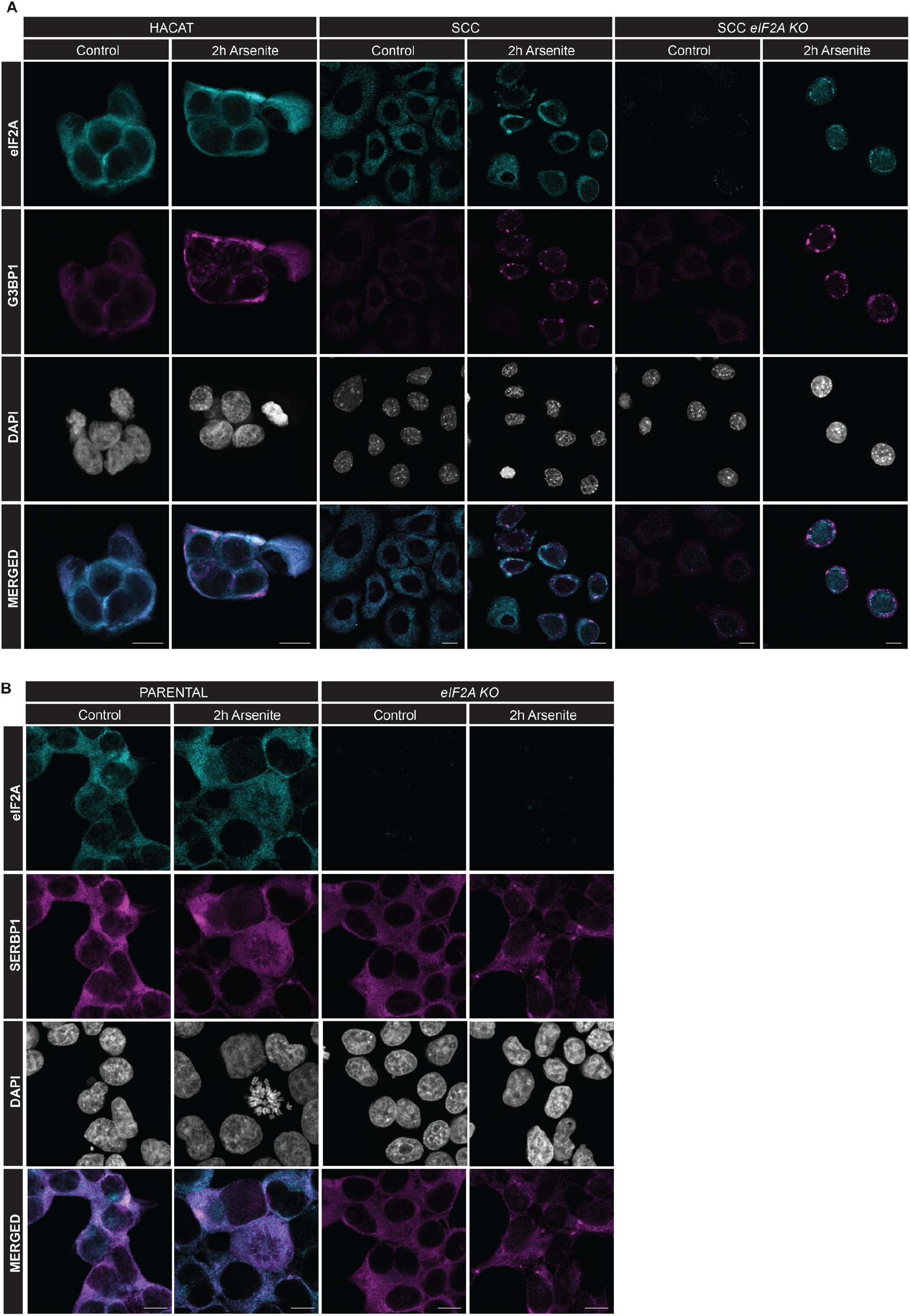
eIF2A localization under stress. **(A)** Confocal microscopy images of eIF2A and G3BP1 in HACAT cells, SCC cells, and *eIF2A knockout* SCC cells following sodium arsenite treatment for 2 hours. Scale bars, 10 µm. **(B)** Confocal microscopy images of eIF2A and SERBP1 in parental and *eIF2A knockout* HEK 293T cells stressed with sodium arsenite for 2 hours. Scale bars, 10µm.

**Figure S7:**
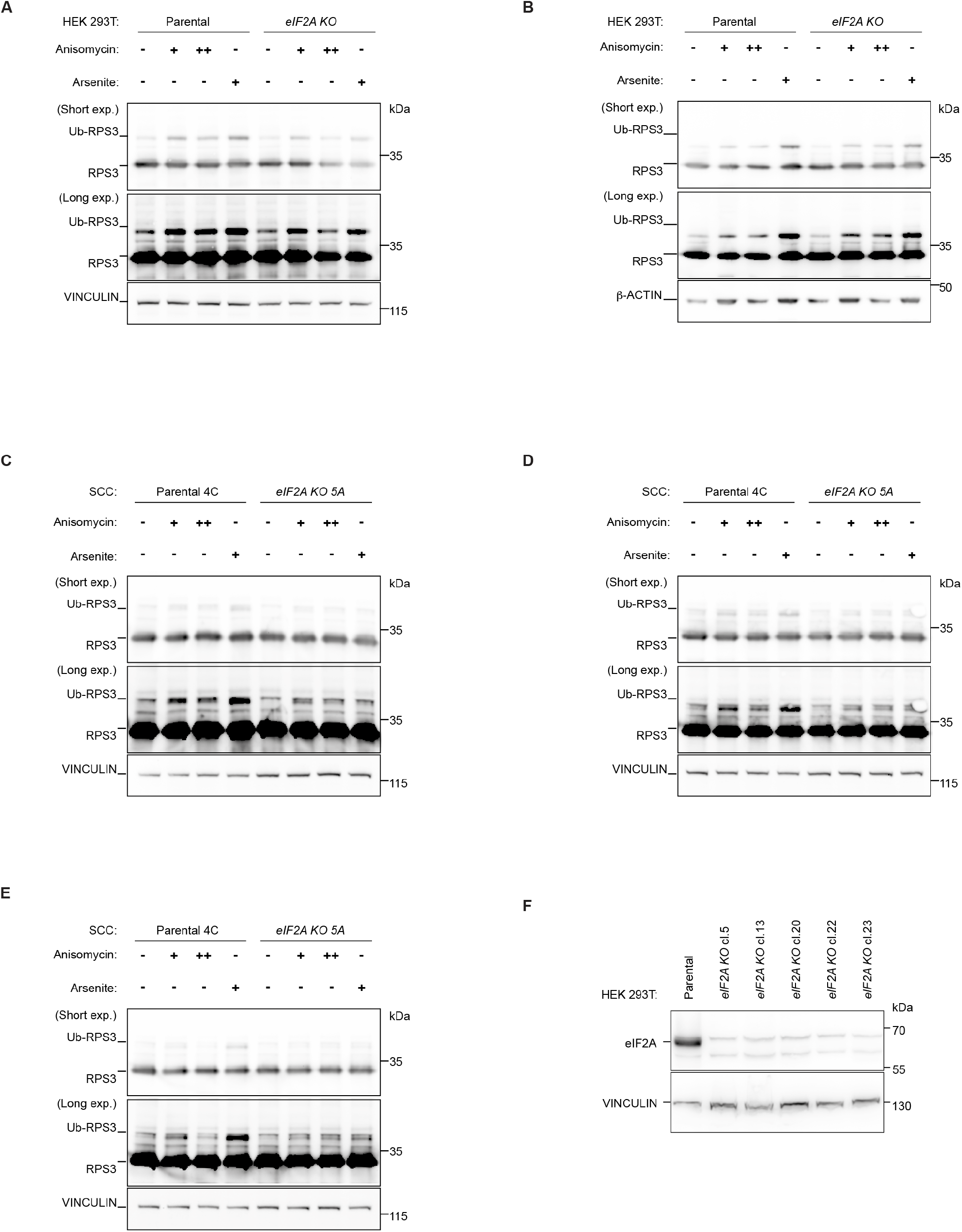
RPS3 ubiquitination upon stress. **(A-B)** Western blot analysis in HEK 293T cells upon anisomycin treatment as individual replicates. Replicate 2 **(A)** and replicate 3 **(B)** alongside replicate 1 (shown in Figure 4A) displaying RPS3 ubiquitination upon anisomycin-induced ribosome collision or stalling. HEK 293T cells were incubated with intermediate dose (+, 1µM) or high dose (++, 100µM) anisomycin for 30 minutes prior to harvesting. In parallel, cells were stressed with sodium arsenite for 2 hours. Quantification of the 3 replicates is displayed in Fig. 4B. Short exp.: shorter exposure. Long exp.: longer exposure. **(C-E)** Western blot analysis upon anisomycin treatment as individual replicates in SCC cells. SCCs were incubated with intermediate dose (+, 1µM) or high dose (++, 100µM) anisomycin for 30 min prior to harvesting. In parallel, cells were stressed with sodium arsenite for 2 hours. Quantification of the Ub-RPS3 bands normalized to the loading controls is displayed in Fig. 4B (n=3). Short exp.: shorter exposure. Long exp.: longer exposure. **(F)** Western blot analysis of different knockout HEK 293T clones generated in this study. Clone 5 was used in all subsequent experiments.

**Figure S8:**
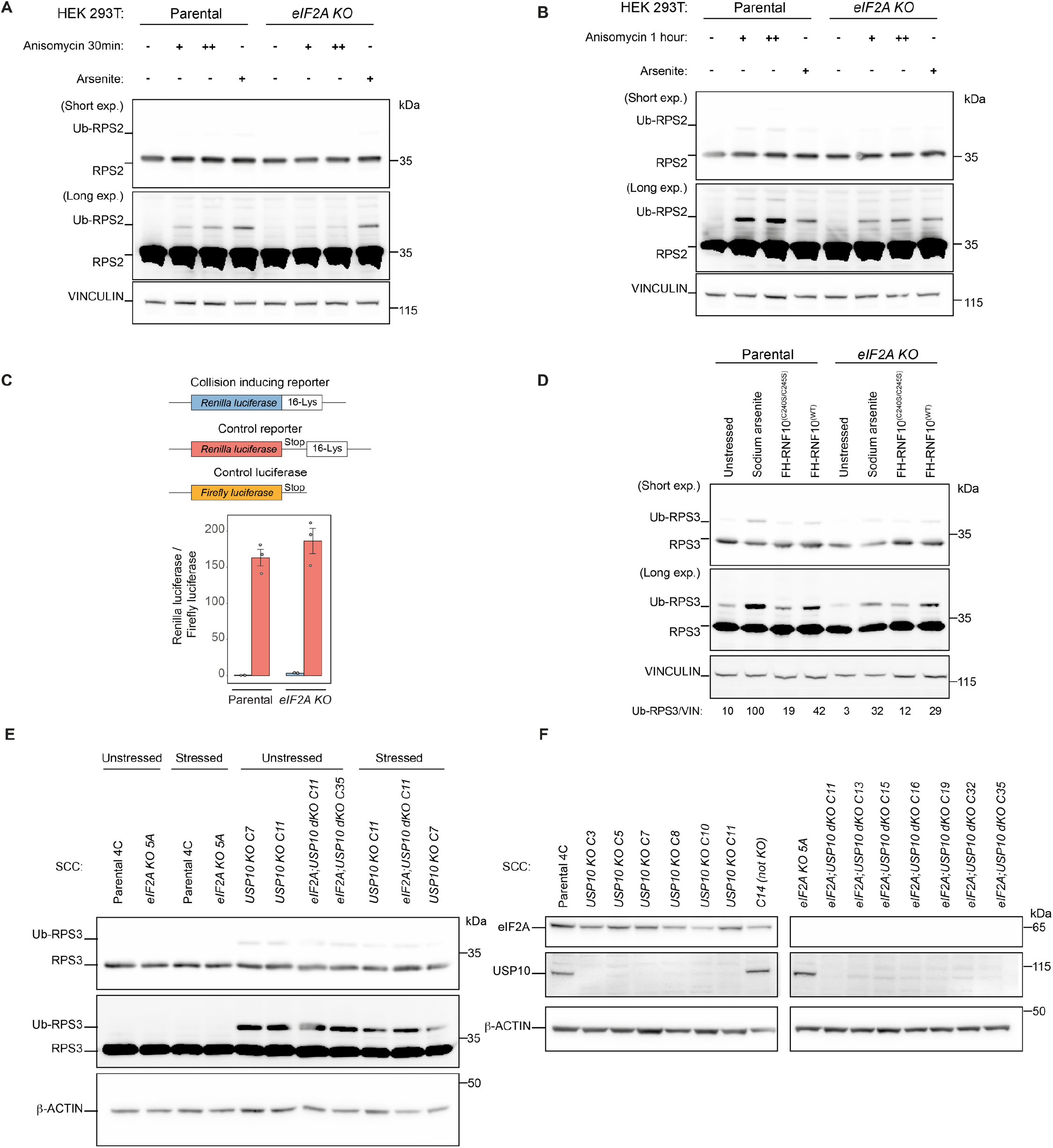
RPS2 ubiquitination and loss of USP10. **(A-B)** RPS2 ubiquitination upon anisomycin treatment. HEK 293T cells were incubated with intermediate dose (+, 1µM) or high dose (++, 100µM) anisomycin for 30 minutes **(A)** or 1 hour **(B)** prior to harvesting. In parallel, cells were stressed with sodium arsenite for 2 hours. Short exp.: shorter exposure. Long exp.: longer exposure. **(C)** Luciferase assays showing the expression of *Renilla* luciferase with a stall-inducing reporter. The expression of the collision-inducing reporter was not significantly increased upon loss of eIF2A. **(D)** RNF10 overexpression induces Ub-RPS3 also in *eIF2A knockout* cells. Western blot images displaying overexpression of human RNF10 constructs (FLAG::HA::RNF10^Wildtype^ and FLAG::HA::RNF10^(C240S/C245S)^ catalytically inactive mutant) in parental and *eIF2A knockout* HEK 293T cells. **(E)** Western blot image showing RPS3 ubiquitination in additional *Usp10 knockout* SCC clones. Cells were stressed with 50 µM sodium arsenite for 2 hours. **(F)** Western blot image confirming loss of USP10 in different *Usp10 knockout* SCC clones.

**Figure S9:**
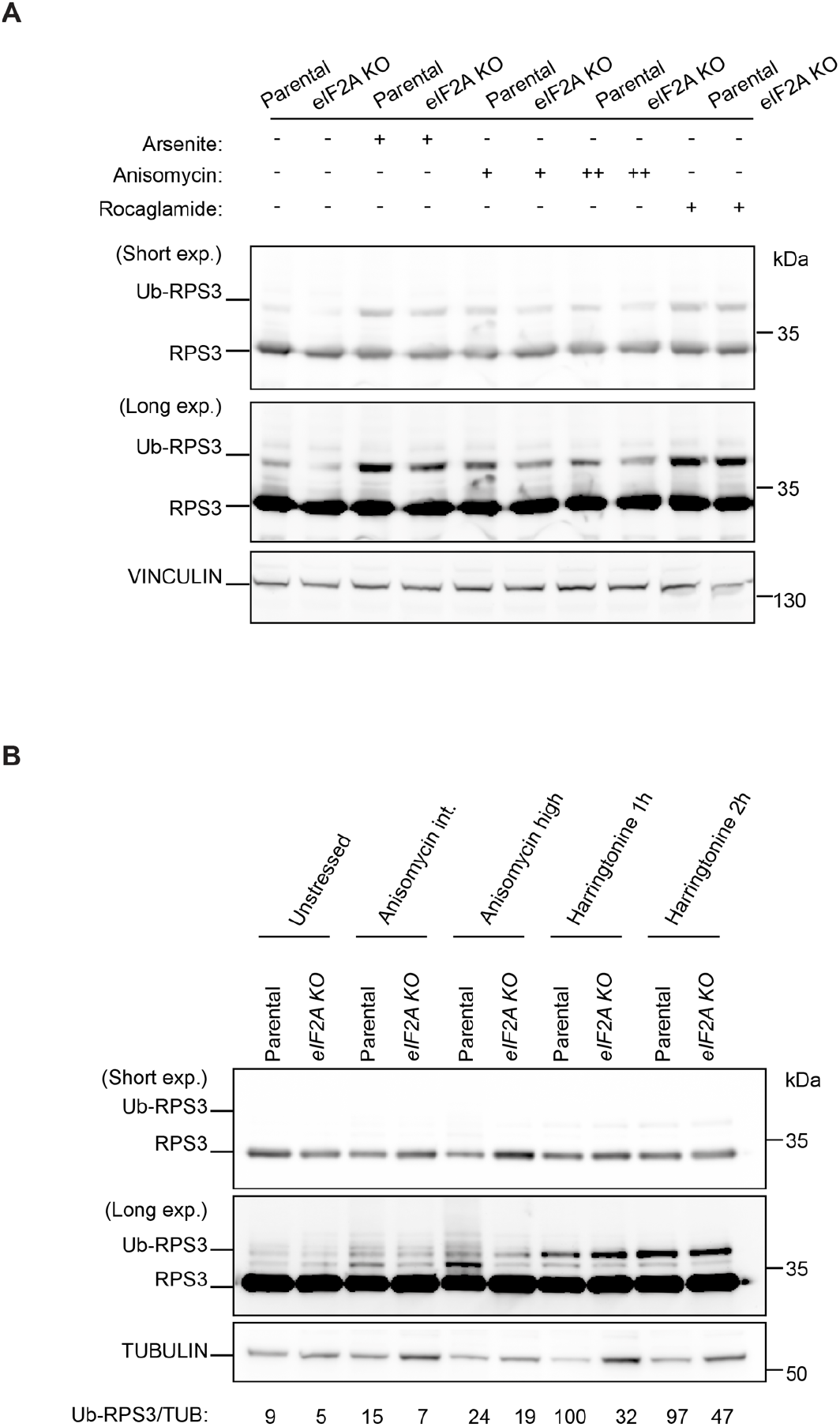
eIF2A does not specifically impact iRQC. **(A)** Western blot analysis shows RPS3 ubiquitination in HEK 293T cells treated with sodium arsenite, intermediate-dose anisomycin, high-dose anisomycin or the eIF4A1 inhibitor rocaglamide, an inducer of iRQC. **(B)** Western blot analysis shows RPS3 ubiquitination in HEK 293T cells treated with intermediate-dose anisomycin, high-dose anisomycin, or the translation initiation inhibitor harringtonine.

**Figure S10:**
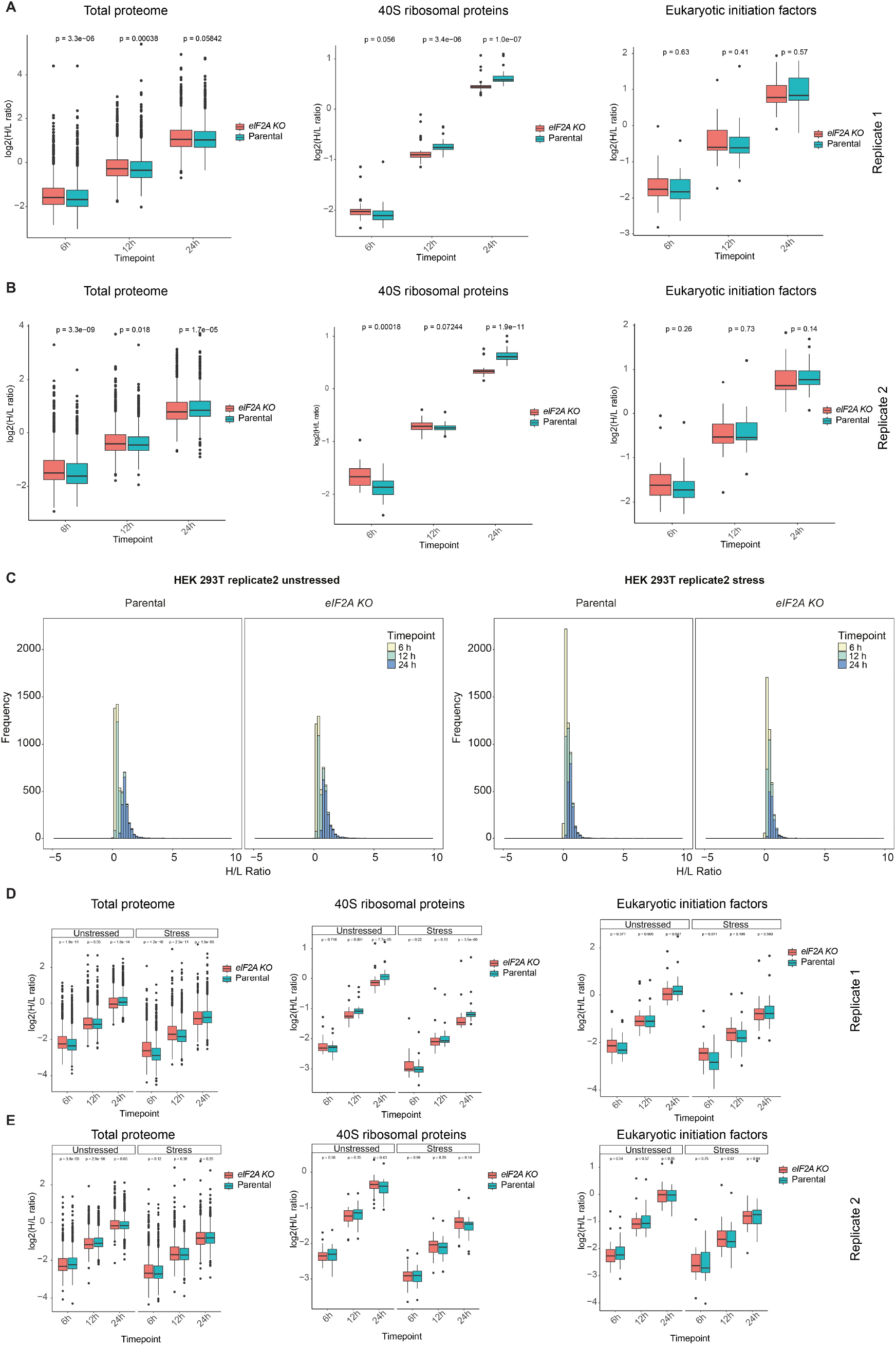
Dynamic SILAC uncovers that eIF2A impacts 40S turnover upon stress. **(A)** Boxplot representation of H/L ratios (replicate 1) across a time course in SCCs, comparing the total proteome, 40S small ribosomal subunit proteins and eukaryotic initiation factors. **(B)** Boxplot representation of H/L ratios (replicate 2) across a time course in SCCs, comparing the total proteome, 40S small ribosomal subunit proteins and eukaryotic initiation factors. **(C)** Histograms displaying the distribution of H/L ratio in parental and *eIF2A knockout* of the replicate 2 across a time course. **(D)** Boxplot representation of H/L ratios (replicate 1) across a time course in HEK 293T, comparing the total proteome, 40S small ribosomal subunit proteins and eukaryotic initiation factors. **(E)** Boxplot representation of H/L ratios (replicate 2) across a time course in HEK 293T, comparing the total proteome, 40S small ribosomal subunit proteins and eukaryotic initiation factors.

